# Unifying mutualism diversity for interpretation and prediction

**DOI:** 10.1101/304295

**Authors:** Feilun Wu, Allison J. Lopatkin, Daniel A. Needs, Charlotte T. Lee, Sayan Mukherjee, Lingchong You

## Abstract

Coarse-grained rules are widely used in chemistry, physics and engineering. In biology, however, such rules are less common and under-appreciated. This gap can be attributed to the difficulty in establishing general rules to encompass the immense diversity and complexity of biological systems. Even when a rule is established, it is often challenging to map it to mechanistic details and to quantify these details. We here address these challenges on a study of mutualism, an essential type of ecological interaction in nature. Using an appropriate level of abstraction, we deduced a general rule that predicts the outcomes of mutualistic systems, including coexistence and productivity. We further developed a standardized calibration procedure to apply the rule to mutualistic systems without the need to fully elucidate or characterize their mechanistic underpinnings. Our approach consistently provides explanatory and predictive power with various simulated and experimental mutualistic systems. Our strategy can pave the way for establishing and implementing other simple rules for biological systems.

---

How populations interact to generate various outcomes is a key question in biology. Mutualism, where two or more populations provide reciprocal benefit, is an essential type of ecological interaction (1). In marine ecosystems, coral reefs are based on mutualistic interactions between coral and algae that provide ecosystem services for humans and habitats for diverse organisms (2). Plant-bacterial mutualism is estimated to generate 60% of the annual terrestrial nitrogen input (3). Cross-feeding in microbial communities is also a mutualistic interaction that influences community structures and is the cornerstone of various microbial metabolic tasks (4, 5). Although mutualistic coexistence is beneficial in maintaining the biodiversity, function and stability of ecosystems, under some conditions mutualistic systems can collapse, where one or more mutualistic partners is lost. This perturbation can further trigger the extinction or invasion of other populations and alter ecosystem functions (6–9). A framework to interpret and predict mutualistic outcomes is useful to prevent undesirable system behaviors and provide guidance for modulating and engineering synthetic mutualistic systems.

Quantitative rules have been developed to elevate our understanding and provide predictive power for various biological systems (10–13). However, such a framework is not yet available for determining mutualism outcomes. Main barriers in developing such a framework are the diversity of mutualistic interaction mechanisms and the complexity of underlying dynamics. Indeed, even engineered mutualistic systems that are by-design capable of cooperation, may not coexist. For example, it is still difficult to predict *a priori* whether an engineered microbial auxotrophic pair can persist or not (14–16). Previously, theoretical criteria in the form of model parameter inequalities have been developed for specific mutualistic systems such as cross-feeding mutualisms (17), plant-pollinator mutualisms (18), seed-dispersal mutualisms (19), ant-plant mutualisms (20), and plant-mycorrhizal mutualisms (21). These criteria depend on the underlying mechanisms *assumed* in the models and are not applicable to other types of mutualistic systems. General criteria have been developed (22–25), such as the classic criterion which states that intraspecific competition must be greater than mutual benefit for a mutualistic system to be stable (26). However, these criteria usually describe transitions between stable coexistence and unbounded growth, and fail to address the transitions between coexistence and collapse and other mutualism dynamics (27, 28) (Fig. S1, Supplementary Text section I).

## Results

### Abstraction reveals a general rule

To reveal any commonality of mutualistic systems, we first summarized the logic of mutualism (Fig. 1a). Mutualism can be defined as the collective action of two or more populations, where each population produces benefit (*β*) that reduces the other’s stress (*δ*) at a cost (*ε*) to itself, *β, δ*, and *ε* are universal features of mutualistic systems (Table 1). In addition to benefit and cost, which are conventionally considered as the driving forces of mutualistic outcomes (29–31), it has been shown that stress factors such as nutrient limitations (32), rising temperature (33, 34), rising CO_2_ levels (35), invasive species (36) are also determining factors of mutualistic outcomes. Although stress is not always explicitly acknowledged in previous models, it has been implemented as decrease in growth rate (37), increase in linear turnover rate (32, 38–41), or increase in intraspecific competition (22, 25). Here we use stress as an umbrella term to capture the effects of stress factors in reducing baseline fitness of individual populations to provide a more complete picture of mutualism (see section II.A of Supplementary Text for the detailed reasoning).

**Figure 1:**
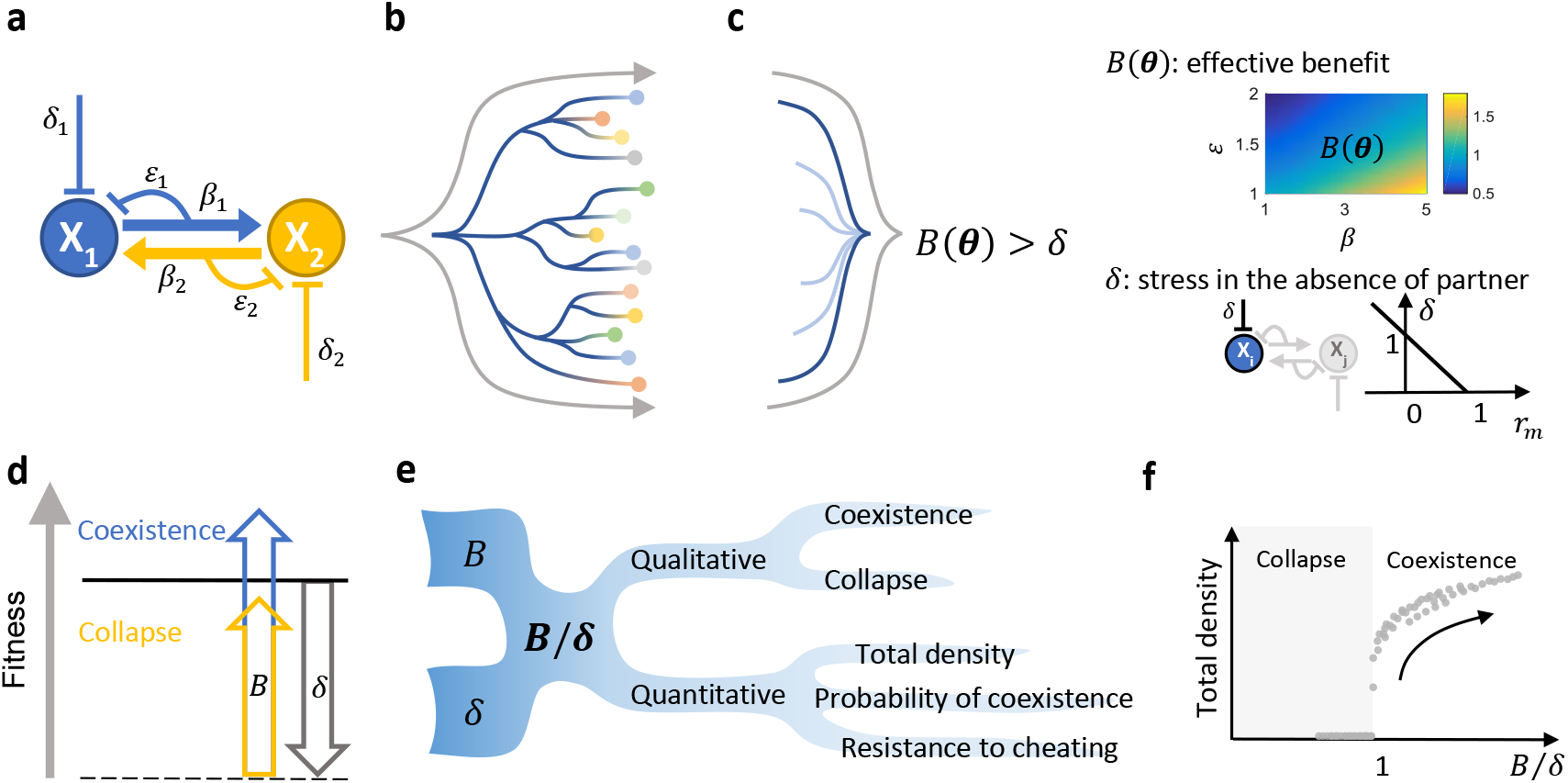
*B* and *δ* are two driving forces that determine qualitative and quantitative mutualistic outcomes **(a) The basic logic of mutualistic systems.** The two partner populations are denoted by *X_1_* and *X_2_*. *β_1_* and *β_2_* describe the strength of benefit the partners provide each other. *ε_1_* and *ε_2_* describe the cooperation cost of providing benefit. Each population also experience stress *δ_1_* and *δ_2_*. **(b) Models originating from the basic logic of mutualism yield diverse coexistence criteria.** Each line represents the generation of a model from the basic mutualism logic and branching represents different implementation details and system complexities. The circles represent the models and different colors represent diverse coexistence criteria derived from these models. This process aims to reflect the diversity of mutualistic systems in nature. **(c) A simple rule emerges at an appropriate level of abstraction.** The lines represent the abstraction process that establishes *Β*(***θ***) *> δ* as the common structure shared by diverse criteria in panel b. *Β*(***θ***) represents effective benefit and is a complex function of model parameters ***θ***, which include *β* and *ε*. In general, *B* increases with increasing *β* and decreasing *ε*. The heatmap is generated using the following model:

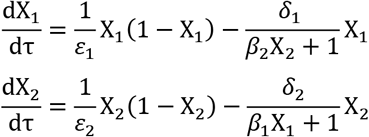

*δ* is the stress experienced by one population in the absence of its partner. **(d) Intuitive interpretation of the simple rule.** The effective benefit *B* must overcome stress *δ* for the system to coexist. Solid black line represents coexistence boundary and dashed black line represents baseline fitness level in the absence of partner. Blue represents a *B* that is greater than *δ* (coexistence) and yellow represents a *B* that is smaller than *δ* (collapse). **(e) *B/δ* can predict various system outcomes.** If the two features *B* and *δ* are known, many downstream predictions, both qualitative and quantitative, can be made. **(f) Quantitative outcomes versus *B/δ***. Simulation results show when *B/δ > 1*, it is predictive of total density. Note that the points do not necessarily lie on a single curve, but a positive trend is well-maintained. Other quantitative outcomes also follow similar positive trends when plotted against *B/δ*.

To reflect the diversity of natural mutualistic systems, we systematically generated a total of 52 ordinary differential equation models based on the basic logic of mutualism with various implementation details (see Materials and Methods and section II.B of Supplementary Text for model assumptions). These implementation details are designed to comprehensively cover the various common and plausible forms of kinetic models that have been adopted in previous studies (see section II.C of Supplementary Text for a summary). Specifically, the models all revolve around the logistic growth equation but differ in the locations of *β, ε*, and *δ* and in the complexities the models capture, such as competition, partner-density-dependent cost and asymmetry (see section II. D of Supplementary Text). We only increased the model complexity to an extent that closed-form steady state solutions are obtainable. We derived coexistence criteria for all 52 models and found that these criteria exhibit diverse structures (Fig. 1b; Table S1–S2; Supplementary MATLAB code). The diversity of our derived criteria is consistent with the diversity of criteria that already exist in the literature (17–21). This diversity highlights the need to have a general criterion, since the appropriate model formulation for a specific system is often unknown *a priori* and its selection can also be non-trivial (42).

Despite the diversity, at an appropriate abstraction level, however, all criteria follow a simple general form (Fig. 1c):

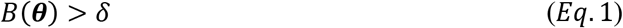

where ***θ*** denotes model parameters accounting for *β* and *ε*. *B*(***θ***) represents the effective benefit produced through mutualistic interaction. Quantitatively, *B*(***θ***) increases with increasing *β* and decreasing *ε* and its structure differs depending on the specific model. *δ* represents stress; regardless of the model structure, it can always be quantified as l – *r_m_*, where *r_m_* is the growth rate of the population in the absence of its mutualistic partner and normalized by its maximum growth rate. The interpretation of our criterion is intuitive: mutualistic partners can coexist if the effective benefit exceeds stress (Fig. 1d). Note that although alternative forms of the criterion may exist, Eq.1 is the most intuitive and parsimonious structure that decouples the effect of mutualistic interaction (*B*(***θ***)) from the baseline stress level of single populations without interaction (ő).

Beyond determining qualitative system outcomes (coexistence versus collapse), *B/δ* defines a general metric that is positively correlated with quantitative mutualistic outcomes (Fig. 1e), such as final population density, probability of coexistence, and resistance to cheater exploitation. This quantitative predictive power of the metric is robustly maintained for both symmetric and asymmetric systems, including obligate and facultative mutualistic systems (Fig. S2a-f; Supplementary Text section IV). Further, the prediction accuracy of our criterion for qualitative outcomes is also robustly maintained in the presence of noise (Fig. S2g). The generality of the metric indicates that it is a property of mutualism. If so, *B* is a high-level feature that, along with *δ*, provides a unifying framework for interpreting and predicting diverse mutualistic systems.

### A calibration approach to use the metric

Quantification of both *B* and *δ* are required to use the metric. Although *δ* is easy to measure by the growth of single populations using a model-independent approach (see Fig. S3 for the general procedure), quantification of *B*, which describes the interactions, can often be challenging. Beyond the difficulties of selecting an appropriate structure of *B*(***θ***), quantification of its underlying parameters often requires nontrivial mechanistic characterizations, such as model parameter fitting and specific biochemical assays. These mechanistic characterizations are especially challenging for cooperative traits, even in well-defined synthetic systems (43–45). Applications of the criterion would thus be difficult for individual systems, let alone enabling streamlined applications for diverse mutualistic systems.

To bypass these challenges, we developed a calibration procedure to use qualitative outcomes to directly quantify *B* as an empirical function of system context ***v***, denoted by *B*(***v***) (Fig. 2a). ***v*** is comprised of system variables that modulate system behaviors, such as temperature, nutrient availability and genetic variation. ***v*** measurements are often readily available, especially in laboratory settings where they are experimentally controlled independent variables. Thus, using simple measurements, we can approximate the true *B*(***θ***(***v***)) that describes the diverse and complex interaction mechanisms without characterizing the specific mechanistic details. *B*(***v***) along with *δ*, will serve as the basis for interpretation and prediction beyond initial data. Although the procedure requires initial measurements of *qualitative* outcomes, *B*(***v***)/*δ* can also provide predictive power for *quantitative* outcomes (Fig. 1e). *B*(***v***)/*δ* is positively related to the final density (Fig. 1f; Fig. S2c-d), probability of coexistence (Fig. S2a, e), and cheater resistance (Fig. S2b, f). Further, *B*(***v***) can be used to reveal how multiple system variables collectively alter the effectiveness of the interaction, which is a major challenge in studying context dependency of mutualistic outcomes (46).

**Figure 2:**
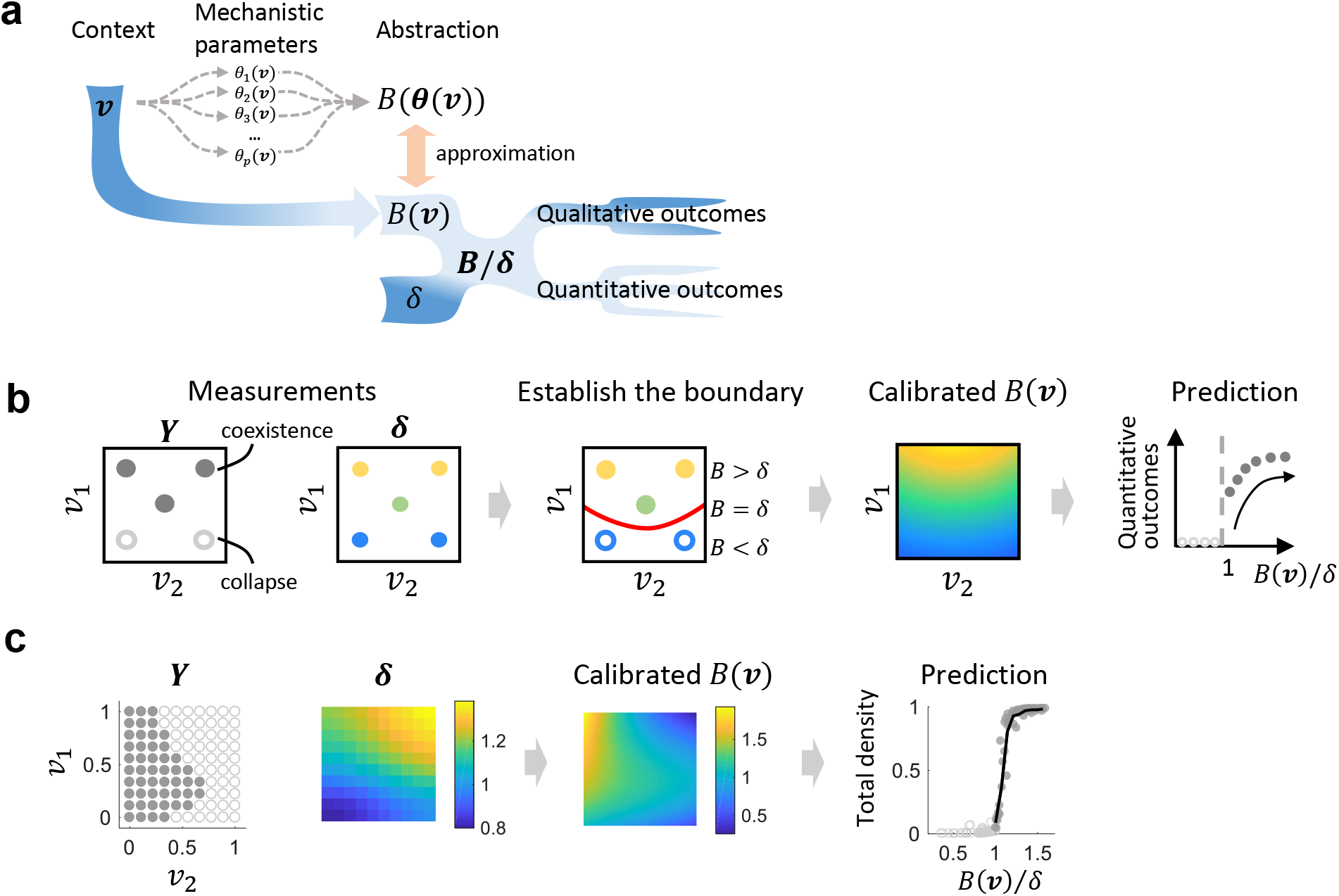
A streamlined approach to calibrate for an empirical *B*(***v***). **(a) The rationale behind the calibration procedure**. Conventional approaches (denoted by dashed gray arrows) require quantifications of mechanistic parameters as functions of contextual variables *θ*(***v***) and finding the appropriate structure of *B*(*θ*) to construct *B*(*θ*(***v***)). However, both steps are challenging and require case-by-case applications due to the diversity of mutualistic interactions. Instead, using qualitative outcomes of the system, we can distill an empirical function *B*(***v***) to approximate the true *B*(*θ*(***v***)). *B*(***v***)/*δ* can then predict qualitative and quantitative outcomes. Dark blue indicates the data that are relatively easy to measure without requiring mechanistic understanding of the interaction. **(b) A schematic demonstrating the mathematical basis of the calibration procedure**. *v_1_* and *v_2_* represent two system variables. A circle represents an observation *i* at a particular ***v_i_*** = (*v_1i_, v_2i_*). 5 observations are shown. ***Y*** contains qualitative system outcomes for each observation. Closed circles indicate coexistence and open circles indicate collapse; the same notation scheme is used for all following figures. ***δ*** contains the measurement of stress for each observation (lighter colors indicate higher values). Using ***v, Y*** and ***δ***, a boundary that separates the two types of outcomes can be established (the red curve). According to our simple rule, *B* = *δ* is true at the boundary; *B > δ* is true for coexistence and *B < δ* is true for collapse. Using these data and our simple rule, we can calibrate for a *B* (***v***) which then enables the interpretation and prediction of system outcomes. Refer to Movie S1 for a 3D visualization of the calibration. **(c) Proof of principle using simulated data**. Simulations were performed using a complex mutualism model that does not have an explicit form of *B*(***θ***) (see Supplementary Text section section V.F). The input data set contains 100 observations. *δ* and calibrated *B*(***v***) share the same axes with ***Y*** (this applies to all following figures). *B*(***v***)/*δ* correctly classifies 97.2% of 2500 new data points. *B*(***v***)/*δ* is also predictive of total densities (only 100 data points are shown out of 2500). Black trace in the plot named “Prediction” represents binned averages of total density (this applies to all following figures).

We first defined the input-output relationship of the calibration procedure (Fig. 2b). Measurements of qualitative outcomes are denoted as ***Y*** = [*y_1_,y_2_,y_3_, … y_n_*] (*y_i_* = 1 for coexistence and –1 for collapse; *i* represents the index of an observation; *n* represents the total number of observations). Measurements of *δ* for the same set of observations are denoted as ***δ*** = [*δ*_1_, *δ*_2_, *δ*_3_,… *δ*_n_]. Note that theoretically, quantification of *δ* for any partner is sufficient. However, choosing the partner with a larger dynamic range of *δ* is preferable since it can contain more information content. System context is denoted by ***v*** = [***v***_1_, ***v***_2_, ***v***_3_,… ***v***_n_], where ***v**_i_* is a vector that contains the values of all system variables for observation i. With inputs ***Y**, **δ*** and ***v***, we can establish a smooth boundary between coexistence and collapse described by *F*(*δ*,***v***) = 0. To ensure *B > δ* for coexistence and *B < δ* for collapse, we constrain *F*(*δ,**v***) > 0 for coexistence and *F*(*δ,**v***) < 0 for collapse. Because *B = δ* is true at the boundary, we can deduce that *F*(*B, **v***) = 0. According to the implicit function theorem, if *F*(*B, **v***) = 0 is continuously differentiable, the output *B*(***v***) is implied. A calibrated *B*(***v***) can then enable downstream interpretation and prediction.

To implement the calibration, we used support vector machine (SVM), a machine-learning algorithm for supervised classification (see Supplementary MATLAB code of the implementation). Assuming continuity of *B*(***v***), we designed kernels that are separable in *δ* and ***v*** to train the function *F*(*δ,v*) = 0. We implemented linear, quadratic, cubic and sigmoidal kernels to describe different possible shapes of *B*(***v***). Because there are infinite number of *B*(***v***) that can provide equivalently high classification accuracy, we ranked the *B*(***v***) obtained from different kernel parameters to find the *B*(***v***) that are closer to the true *B*(***θ***(***v***)) (Fig. S4a). The ranking method is established using simulated data where the true *B*(***θ***(***v***)) is known, so that each *B*(***v***) can be evaluated against it by coefficient of determination (R^2^). We found that our procedure consistently optimizes for R^2^ (Fig. S4b-c; Supplementary Text section V). The proper sample size for the calibration can be evaluated based on the exponential decay of bias with increasing sample size (47) (Fig. S4d).

Using this procedure, we first tested whether *B*(***v***)/*δ* can be applied to mutualism models in which no explicit form exists for *B*(***θ***). To do so, we constructed an overwhelmingly complex model with competition, partner-density-dependent cost, high Hill coefficient and asymmetric function structures (see Supplementary Text section V.F, Fig S5a). Using an input data set of 100 points, *B*(***v***)/*δ* correctly predicts coexistence versus collapse for 97.2% of test data beyond the initial 100 data points. As expected, *B*(***v***)/*δ* provides predictive power for quantitative outcomes including total population size (Fig. 2c), probability of coexistence (Fig. S5b) and resistance to cheater exploitation (Fig. S5c). The confidence of *B*(***v***) can be evaluated by the consistency of top *B*(***v***) and their relative standard deviation (RSD) (Fig. 5 d, e).

### Experimental application of the metric

We next applied our framework to three experimental systems to test its applicability. As the first example, we engineered two synthetic mutualistic partners in Top10F’ strain of *Escherichia coli*, denoted by M_1_ and M_2_ (Fig. 3a, Fig. S6a). In this system, stress is modulated by the concentration of Isopropyl β-D-1-thiogalactopyranoside (IPTG) which induces the expression of CcdB (a toxin). Anhydrotetracycline (aTc) induces quorum sensing (QS) modules in both strains to each produce a unique QS signal that triggers the production of CcdA (the antitoxin of CcdB) in the partner population. The production of aTc-induced expression of the QS module can impose cooperation cost to both strains. Consistent with circuit design, our experimental results demonstrated IPTG-mediated growth suppression and aTc-mediated mutual rescue (Fig. S6b).

**Figure 3:**
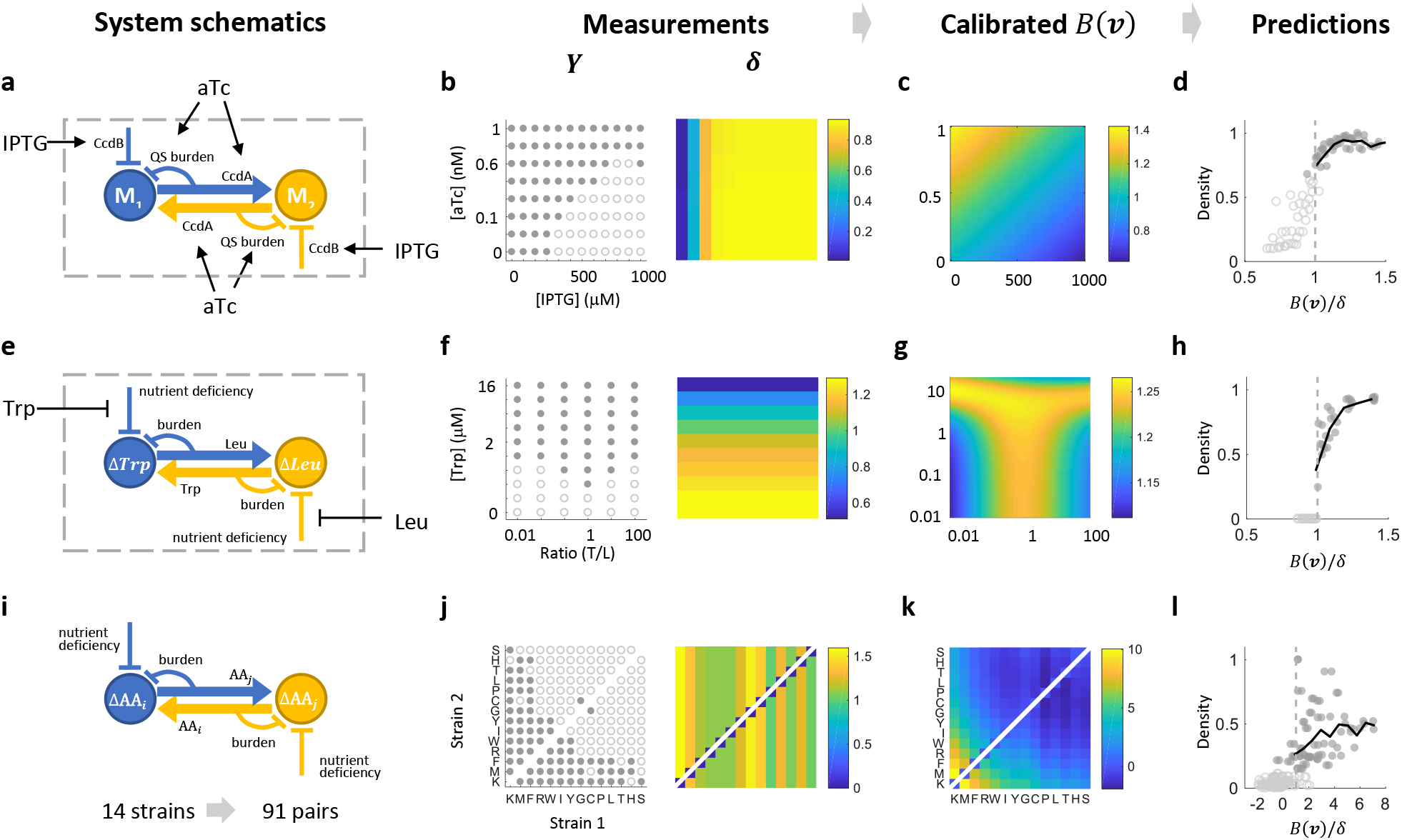
Implementation of the simple rule in experimental systems (Movie S2-4 for 3D visualizations) **(a) The QS-based mutualism system.** IPTG modulates stress and aTc induces QS-mediated mutualistic interaction. **(b) Measurements of coexistence and collapse and corresponding *δ* values**. Coexistence and collapse are measured by co-culturing the two strains starting from the same initial densities. *δ* is measured by OD of M2 mono-culture after 32 hours of culturing. **(c) Empirical calibration of *B*(***v***)**. *B*(***v***) reveals how [IPTG] and [aTc] together modulate the effectiveness of the interaction. **(d) *B*(***v***) /*δ* is predictive of coexistence versus collapse and total final density**. The x-axis range of [0.5, 1.5] is used to highlight the transition (this also applies to other prediction plots). The trend continues to hold beyond this range. The y axis represents normalized final cell density (the same applies to panel h and l). **(e) The pair-wise yeast auxotroph system**. The growth of both auxotrophs are suppressed in monocultures. With increasing Trp and Leu supplemented to the co-culture, the growth suppression can be alleviated. **(f) The amount of supplemented amino acid and ratio of initial densities modulate system behavior**. Only [Trp] is shown and [Leu] is 8 times of [Trp]. The total initial density of the two strains are kept constant. Corresponding *δ* values are measured based on growth yield of *ΔLeu* monocultures, assuming *δ* is independent of initial density (Fig. S7a). **(g) Optimal effective benefit occurs at an intermediate ratio of initial density**. The calibrated ***B***(***v***) provides a cross validation accuracy of 95.0%. **(h) *B*(***v***)/*δ* is predictive of normalized total cell number per culture well**. **(i) The 91 mutualism systems constructed by 14 engineered *E. coli* auxotrophs**. Growth suppression is evident in their inability to survive individually in minimal medium. However, two auxotrophs can potentially survive through mutualistic interaction in a coculture by exchanging amino acids. The metabolic burden of the exchanging amino acid can be minimal. **(j) System outcomes for all 91 pairs and *δ* for each of the 14 auxotrophs**. Note that for one pair, the calibration is done twice with *δ* of either strain. **(k) The predictive power of *B*(***v***)/*δ***. The calibrated *B*(***v***) and *δ* measurements provide a 91.8% cross validation accuracy. **(l)** *B*(***v***)/*δ* is predictive of the normalized fold change of final total density relative to initial density.

We cocultured the two strains starting from the same initial density with different concentrations of IPTG and aTc, which are the two dimensions of ***v***. The outcomes of coexistence and collapse are evident in the bimodal distribution of optical density (OD) at 32 hours of culturing (Fig. S6c). *δ* can be quantified by treating monocultures with the same set of [IPTG] and [aTc]. We used *δ* for M2 since it has a wider dynamic range (Fig. S6d). Using these data (Fig. 3b), we obtained a calibrated *B*(***v***). The confidence of *B*(***v***) is evaluated by the consistency of the top 5 *B*(***v***) and relative standard deviation of each *B*(***v***) (Fig. S6e). Consistent with the circuit logic, *B*(***v***) increases with increasing [aTc]. The calibration reveals that [IPTG] also modulates *B*(***v***), which indicates unintended system complexities, such as QS cross-talk and unequal fitness of the two populations (Fig. 3c). We used cross-validation to evaluate how well new observations can be predicted. We found that *B*(***v***)/*δ* provides a cross validation accuracy of 96.8% for coexistence versus collapse and it is also predictive of total final density (Fig. 3d).

We then applied our procedure to data on a pair of *Saccharomyces cerevisiae* auxotrophs that is previously published (32). In this system, one strain cannot produce tryptophan (Trp) and the other cannot produce leucine (Leu). The mutualistic interaction of this system is realized by the exchange of the two amino acids in cocultures (Fig. 3e). Because [Leu] is maintained as 8 times of [Trp], we use [Trp] as one of the dimensions of ***v*** to represent overall concentration of supplemented amino acid. The ratio of initial densities of the two strains composes the other dimension of ***v*** (Fig. 3f, Fig. S7a). All top 5 *B*(***v***) reveal that intermediate ratios of initial density and increasing amino acid concentrations elevate *B*(***v***) (Fig. 3g). However, at the highest level of supplemented amino acid ([Trp] = 16nM, [Leu] = 128nM), top-ranked *B*(***v***) have qualitatively different trends, indicating a low confidence of *B*(***v***) at high concentration (Fig. S7b). Interestingly, this high variability coincides with the system transitioning into competitive exclusion (32). Nevertheless, *B*(***v***)/*δ* is still predictive of final densities (Fig. 3h). Furthermore, we explored using the concentration of supplemented amino acid as a single system variable. *B(**v**)/δ* in this case can also predict the probability of coexistence as the ratio of initial densities varied (Fig. S7c).

In the third example, we applied our framework to previously published measurements of 14 engineered auxotrophic *E. coli* strains that compose 91 pairwise mutualistic systems (48) (Fig. 3i). The genetic context of the two partners vary while the growth environment is kept the same. The classification of coexistence versus collapse is based on the bimodal distribution of total density (Fig. 8a). *δ* of each auxotroph is determined based on final cell densities of monocultures when supplemented with different concentrations of its corresponding amino acid (Fig. S8b). We sorted the auxotrophs by the number of partners they coexist with to convert categorical indices into an ordinal scale. Thus, ***v*** is composed of ordinal rankings of the two strains and measurements of coexistence versus collapse and *δ* are both arranged accordingly (Fig. 3j). We used strain 1 as the reference strain for the calibration. The calibrated *B*(***v***) generated a cross validation accuracy of 91.8% and we verified that *B(**v**)/δ* is predictive of final total density (Fig. 3k-l, Fig. S8c). We noticed a relatively high level of variability of total density when *B(**v**)/δ* > 1, which can be due to system-specific properties that are not fully accounted for by mutualistic interactions.

### Generalization of the framework

In nature, mutualism can occur among three or more partners (49). Thus, we tested our framework with simulations and experimental measurements of N-mutualist systems. Here we show the calibration procedure with simulated data from a 5-mutualist system and found that the quality of the calibration results is well-maintained (Fig. 4a; Fig. S9a; Supplementary Text section VI.A). The same 14 auxotrophs (48) presented above were also used to construct all possible three-member systems. Using the same procedure with a 3-dimensional ***v***, where each dimension represents the genetic context of one strain, we found the predictor *B(**v**)/δ* provides an 89.3% cross validation accuracy and remains predictive of the total density, which indicates the scalability of the framework in experimental settings (Fig. 4b, Supplementary Text section VI.B). We hypothesized that *B*(***v***) calibrated for pairwise interactions can be used to directly construct a metric for 3-member systems, since theoretical analysis shows that n-member *B(**θ**)* can be approximated by pairwise *B(**θ**)* (Table S2). We assume *B* of a 3-member system is the average of *B* for all 3 combinations of its underlying 2-population systems and the same is true for *δ*. The constructed *B/δ* for 3-member systems explains 80.8% of system outcomes (Fig. S9b). This result suggests the possibility of directly extending *B* and *δ* from simple systems to more complex systems without further calibration.

**Figure 4:**
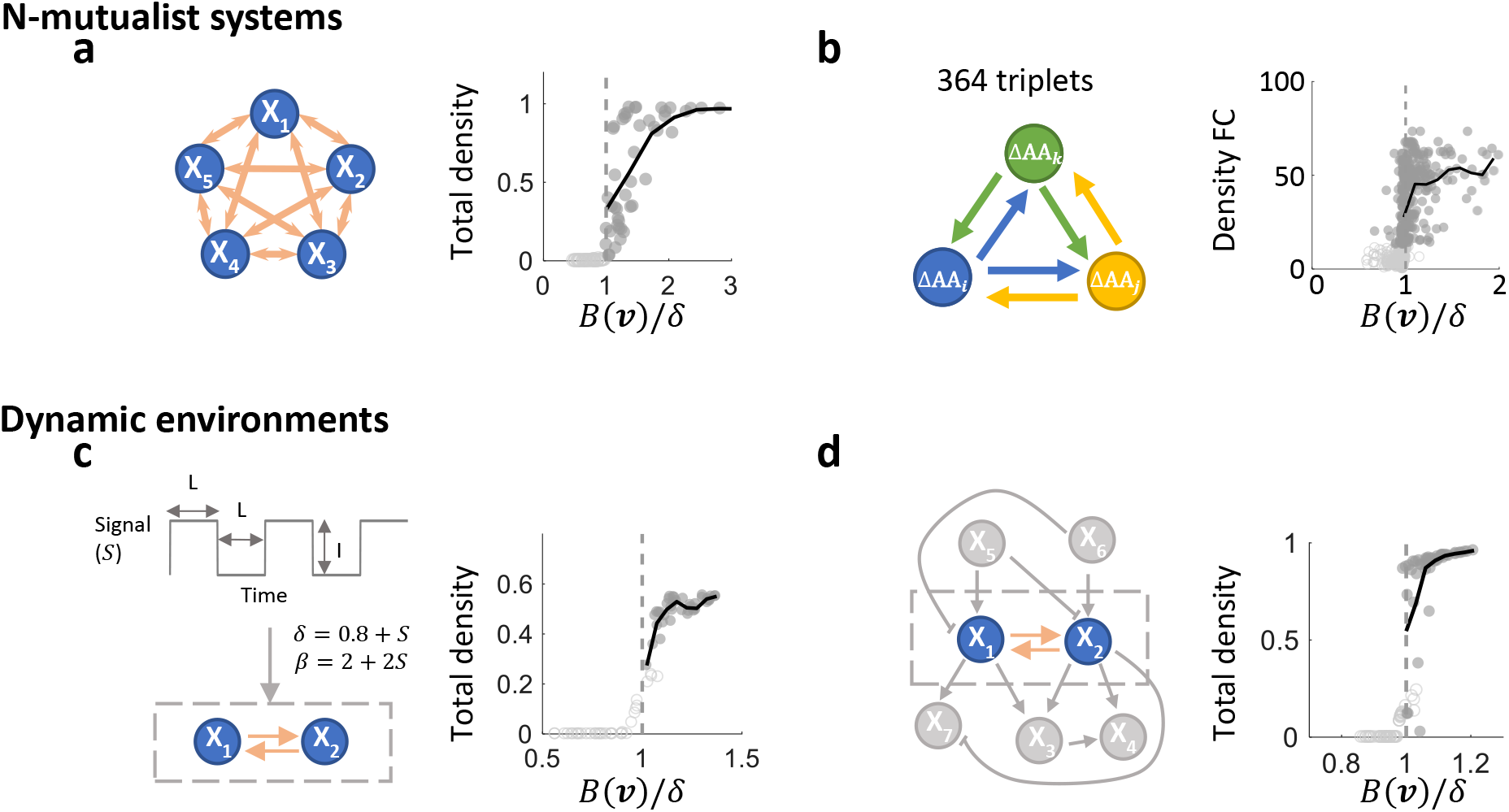
Our framework is generalizable to mutualistic systems with more than two partners and systems that inhabit dynamic environments. Please refer to section VI of Supplementary Text for simulation details. **(a) A simulated mutualistic system with 5 partners.** Parameters in the model are functions of two independent variables. Using 100 data points, we obtained a *B*(***v***) through clibration. *B(**v**)/δ* successfully predicts coexistence versus collapse for 98.6% of a new set of 2500 data points (100 points are shown) and it is also predictive of total density. **(b) Experimental auxotrophic triplets that are comprised of 14 *E. coli* auxotrophs**. The same auxotrophic strains presented in Fig. 3i–k were used to construct three-way mutualistic systems. This experimental validation demonstrates the generality of framework beyond pairwise interactions. **(c) A system that is modulated by an oscillatory signal**. The oscillatory signal *S* is described by *I* (intensity) and *L* (duration). The signal temporally modulates *δ* and *β. I* and *L* are the two system variables used in calibration. The procedure achieves a prediction accuracy of 97.3% for new data. **(d)** A simulated mutualistic system that coinhabits with 5 other populations. X_1_ and X_2_ are the mutualistic partners. X_3_ to X_7_ are bystander populations that either modulate or are modulated by X_1_ and X_2_. *B(**v**)/δ* successfully predicts 92.3% of new data in this example.

Beside static environments, mutualistic systems can also inhabit dynamic environments where they experience fluctuating physical and chemical cues or cohabitate with other populations. We verified that the theoretical criterion generally holds in both cases (Fig. S10a). However, the transition between collapse and coexistence does not strictly occur at 1, which further advocates for the necessity of the calibration process. With simulated data, we carried out the calibration procedure and verified that the applicability of our framework is well-maintained (Fig. 4c-d, Fig. S10b, Supplementary Text section VI.C-D). The robustness of the framework suggests that it can be used to study microbial communities, of which advancements in both interpretation and prediction are in demand (50).

## Discussion

The immense complexity and diversity of biological systems is intriguing and inspires the exploration of mechanistic details. However, these details can distract us from simple rules that emerge at a higher level. By abstracting away from low-level details, many simple rules for biological systems have been developed to enhance our understanding and provide predictive power (51, 52). A classic example is the Hamilton’s rule, which states that a cooperative trait will persist if *c/b > r*, where *r* is the relatedness of the recipient and the actor; *b* is the benefit gained by the recipient; and *c* is the cost to the actor. More recent examples include linear correlations underlying cell-size homeostasis in bacteria (53–55), ranking of quorum sensing modules according to their sensing potential (56, 57), and the growth laws resulting from dynamic partitioning of intracellular resources (58, 59). Beyond establishing another simple rule, we also demonstrated that one can purposefully seek an appropriate abstraction level where a simple unifying rule emerges over system diversity. If this rule anchors in the basic definition of a type of system, it can then be applied to diverse systems of the same type. Beyond microbial systems that we tested, our criterion in principle can also be applied to other systems of larger or smaller scales that share the same logic.

Although simple general rules in biology are powerful tools, their applicability to experimental systems can be limited by the difficulties in associating the abstracted parameters to lower-level mechanistic details and quantifying these details experimentally. This is evident in the application of Hamilton’s rule to, for example, to experimental systems (43–45). For simple rules that are dictated by inequalities, our calibration procedure provides a tool to apply these rules directly to experimental systems. If one side of the inequality and some final outcomes can be measured, the other side can be calibrated as an empirical function. Although the procedure cannot further dissect the empirical function into specific mechanistic parameters, the function can serve as an overall summary of the underlying mechanistic details while bypassing the requirement of characterizing them individually. Our approach thus enables the downstream interpretation and prediction by these simple rules with readily-accessible measurements.

## Acknowledgements

We thank Tim Hoek and Jeff Gore for providing data used for analysis in Figure 3f-h and Michael Mee and Harris Wang for sharing the data used for analysis in Figure 3j-l. We also thank Yu Tanouchi, Lawrence David, Wenying Shou, Nan Luo, Yangxiaolu Cao, Carolyn Zhang, Ryan Tsoi, Teng Wang, Shangying Wang for constructive inputs.

## Author contributions

F.W. conceived the research, designed and performed both modelling and experiments, interpreted the results, and wrote the manuscript. A.J.L. constructed the synthetic circuit, assisted with experimental design and manuscript revisions. D.N. assisted with modeling, results interpretation, and manuscript revisions. C.L. assisted with establishing the general relevance of the criterion and manuscript revisions. S.M. assisted with establishing the calibration process and manuscript revisions. L.Y. conceived the research, assisted in research design, data interpretation, and wrote the manuscript. All authors approved the manuscript.

## Data and materials availability

All data is available in the manuscript or the supplementary materials.

**Table 1:**
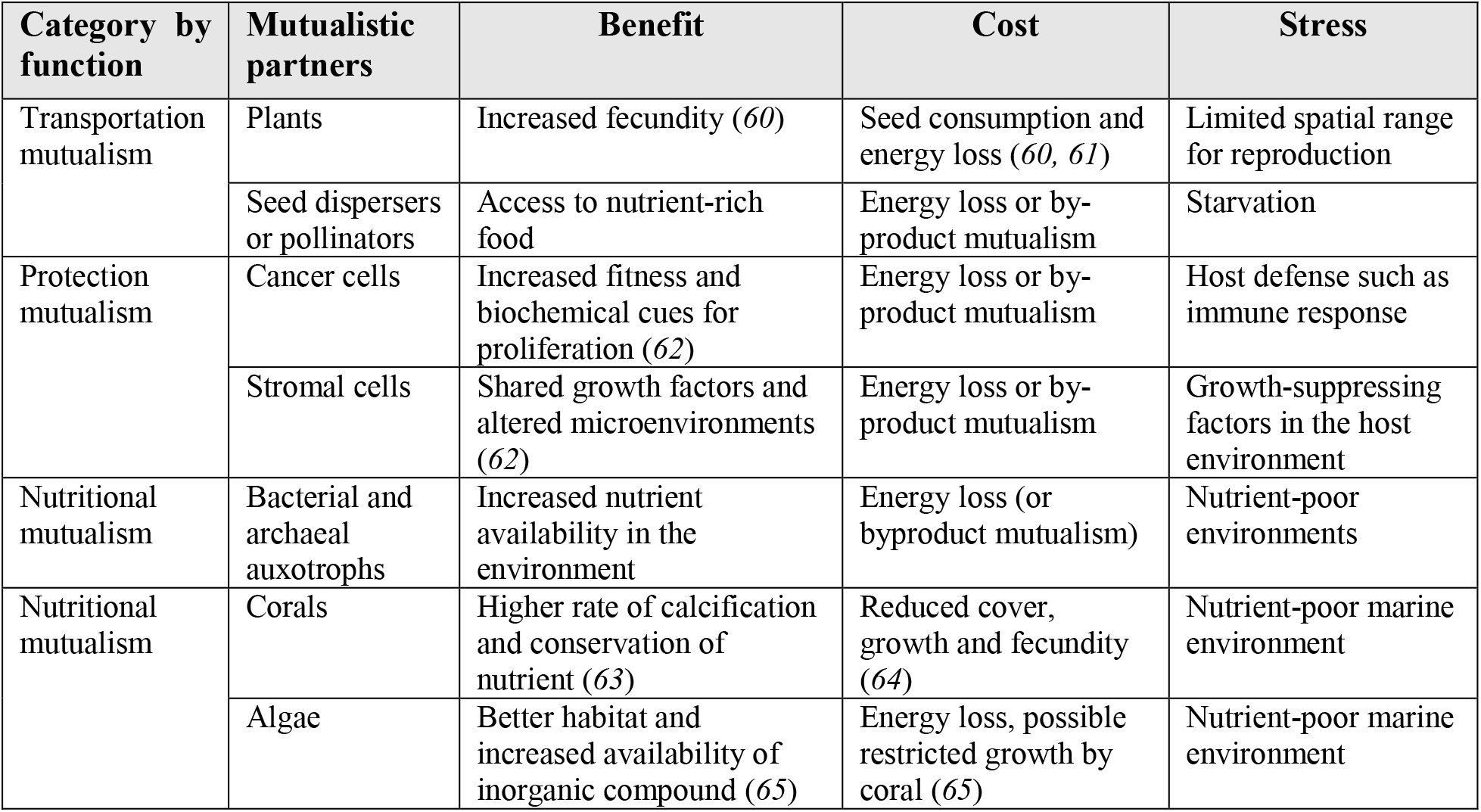
Examples of benefit, cost, and stress in diverse mutualistic systems

## Materials and Methods

### Model development

We built mutualism models based on four key assumptions:

a. Benefit shall increase growth rate or carrying capacity and is positively dependent on partner density.
b. Cost shall *decrease* growth rate or carrying capacity.
c. Stress shall produce negative growth of populations at some parameter combinations.
d. Negative growth of a population shall be potentially counteracted by benefit provided by a partner, but further strengthened by cost.

See SI section II for detailed reasoning and implementation of each assumption.

### Criteria derivation

We calculated the analytical solutions of fixed points of the 52 models using MATLAB R2017a. Then we identified the fixed points that represent stable coexistence. The coexistence criteria are derived by ensuring the fixed points are real positive numbers. We can then rearrange the inequality to have *δ* on one side. The other side of the inequality is then an expression of other parameters, which is expressed as *B(**θ**)*. More details are presented in SI section III. The MATLAB code of the models and the derivation and testing process is included in the Supplementary Materials.

### Calibration of benefit landscape using SVM

We used support vector machine (SVM) algorithms in MATLAB to implement the calibration. The input data are formulated into the following format:

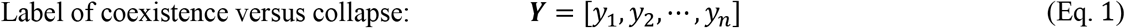

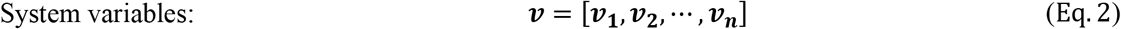

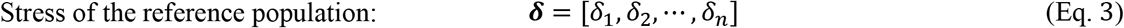

In Eq. 2–4, *n* represents total number of observations and each index represents one observation. *Y* takes values of 1 or – 1, which represent coexistence versus collapse for each observation. ***v*** contains the coordinates in the independent variable space where observations are obtained. ***δ*** contains the stress levels of the reference population for each observation.

We designed kernels that have additive separability between ***v*** and ***δ***, which can be expressed in a general form:

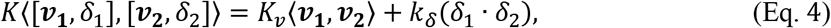

where *K_v_* is the kernel that dictates the shape of the empirical function of *B* and *k_δ_* is a constant that varies the weight of ***δ***. The predictor trained using SVM can then be expressed as:

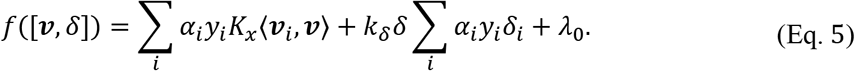

According to our criterion, we know that *B = δ* when *f*([***v***, *δ*]) = 0. We can then derive from Eq. 6 a function of *B* in terms of system variables ***v***:

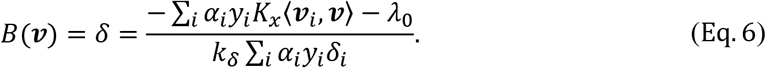

With each set of inputs, we used linear, quadratic, cubic, and sigmoidal kernels with different kernel parameters to train *B*(***v***). We then find the *B*(***v***) with lowest cross-validation classification loss and bootstrapped variance. The top ranked *B*(***v***) is then used along with *δ* measurements for interpretation and prediction. See section V of SI for detailed calibration method and the Supplementary Materials for the MATLAB code of the calibration process and sample data sets.

### QS-based mutualism strains

The two strains were constructed based on circuit components from a synthetic predator-prey system *(66, 67)*. Both populations carry two plasmids. Briefly, M1 carries plasmids identical to the predator plasmids, denoted A1 for the module carrying *ccdA (tet* promoter (68) driving *luxR* and *lasI* followed by *lux* promoter driving ccdA) and B1 for the module carrying *ccdB (Lac* promoter (68) upstream of *ccdB* followed by *tet* promoter upstream of *gfp)*. To construct M_2_, A1 was used as backbone. To obtain orthogonal communication, KpnI and NotI restriction digest cloning was used to replace *luxR/lasI* genes from A1 with *lasR/luxI* genes from the previously published prey plasmid (consisting of pLac lasRluxI CcdB (Kan^R^, p15A ori)). Reporter plasmid B1 is from (66). To construct B2, enzymes XhoI and KpnI were used to replace the *tet* promoter on prey plasmid with the *ccdB* module from B1. All M1 and M2 plasmids were verified using restriction digest and sequencing.

### Growth conditions of QS-based synthetic system

The experiments of QS-based mutualistic system were done in 96-well microtiter plates. PH-buffered M9 medium (M9 salt supplemented with 1mM thiamine, 0.2% casamino acid, 0.4% glucose, 2mM MgSO_4_, 0.1mM CaCl_2_ and buffered with 100mM MOPS with PH adjusted to 7.0) was used. 50 μg/mL kanamycin and 100 μg/mL chloramphenicol were added to the culture to maintain plasmids.

To measure circuit function, 4 ml LB media in a 14 ml culture tube was inoculated from single colony and incubated overnight at 37 °C at 250 r.p.m. The optical density is adjusted to 0.5 in M9 media (measured at 600 nm with TECAN microplate reader) before use. Cocultures are created by mixing both strains in a 1:1 volume ratio. The culture is then diluted 10^6^ fold and cultured in 200 μL batch culture at 30°C in TECAN plate reader to record OD for 32 hours with 10 minutes between each reading. The inducers were added to the media at the beginning with cell culture.

## Supplementary Text

Contents

I. Previous mutualism models
  A. The Lotka-Volterra model
  B. Other variants
II. Building various mutualism models
  A. Incorporation of stress
  B. Model assumptions
  C. xModifications of the logistic growth equation
  D. Specific implementations of benefit, stress and cost
III. Derivation of diverse criteria
  A. An example
  B. Summary of criteria derived from symmetric models
  C. Criteria from models with additional complexities
IV. Theoretical generality of the simple metric
  A. Establish the general predictive power of ***B/δ***
  B. The predictive accuracy is maintained in the presence of noise
V. Calibration using SVM
  A. Kernels
  B. Standardizing input data
  C. From SVM output to calibrated ***B(v)***
  D. Cross validation and bootstrapping
  E. Twenty sets of simulations to establish and test the calibration procedure
  F. Calibration with simulated data with unknown ***B(θ)***: an example
VI. Framework generality verified by complex mutualistic systems
  A. A 5-mutualist system
  B. The experimental 3-member mutualistic systems
  C. A mutualistic system in an oscillatory environment
  D. A mutualistic system cohabiting with other populations

References

## I. Previous mutualism models

Here we briefly summarize some limitations of previously published models in studying the transition between mutualism coexistence and collapse.

### A. The Lotka-Volterra model

Consider the following model that follows the basic form of Lotka-Volterra model of mutualism. This formulation is a nondimensionalized form of previous models (69–74). The parameters are renamed according to our parameter assignments.

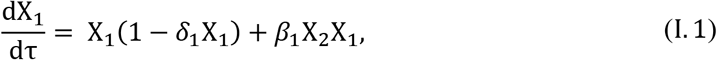

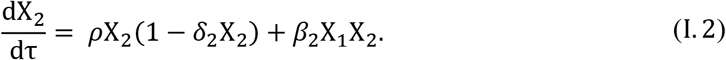

Although the model formulation captures the logic of mutualism, it can generate unbounded growth of the two partners, which is not a biologically relevant state. For example, when *ρ* = 1, *β_1_* = *β_2_* = 2 and *δ_1_* = *δ_2_* = 1, the two populations are not bounded by their carrying capacities, but both grow exponentially. Importantly, the model does not capture population collapse so it cannot explain the transition between collapse and coexistence. Thus, this model formulation is not suitable for our purpose.

A coexistence criterion for mutualism can be derived using L-V model formulation and is previously demonstrated (69–73):

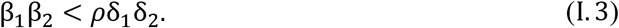

However, this criterion captures the transition between stable coexistence and unbounded growth. It also suggests that mutualism is destabilizing, since increasing the strength of mutualistic interaction (β_1_β_2_) tends to violate the above condition (I. 3). The interpretation of this criterion can be contradictory to empirical observations that mutualism stabilizes community (75–77). Although Lotka-Volterra models are sufficient to answer many questions related to mutualistic systems, it has been proposed that this discrepancy between model dynamics and empirical observations can be attributed to its unrealistic assumptions (78–81).

### B. Other variants

We found that general mutualism models that generate unbounded growth usually do not capture the transition between coexistence and collapse. For example, the following three models were established previously to find structurally stable mutualistic models (78). In contrast to the L-V model which implements the interaction as a linear term, the following models present three alternative ways of adding the interaction to the basic logistic growth equation.

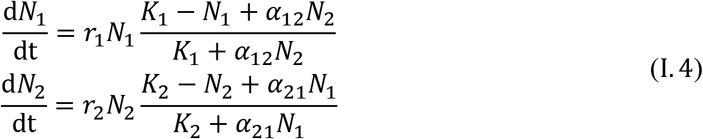

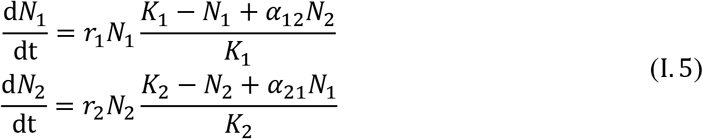

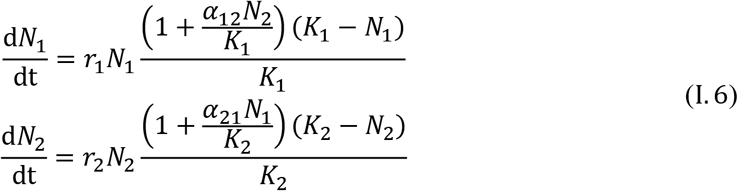

In all these three cases, some parameter combinations generate unbounded growth. For example, *r_1_ = r_2_ = 1; K_1_ = K_2_ = 1; a_12_ = a_21_ = 2; N_10_ = N_20_* = 0.1 generates unbounded growth for I.4 and I.5 and *r_1_ = r_2_ = —1; K_1_ = K_2_ = 1; a_12_ = a_21_ = 2; N_10_ = N_20_ = 2* generates unbounded growth for I.6. Unbounded growth has been recognized as a limitation of many mutualism models (82–84). In addition, although population collapse can be simulated by *r_1_ <* 0 and/or *r_2_ <* 0, in all cases, the fitness of the benefit-receiver decreases with increasing partner density, which is contradictory to the basic logic of mutualism.

To avoid generating unbounded growth, one strategy is to introduce saturating benefit (73). However, although preventing unbounded growth, these models may still not generate negative growth which can be potentially countered by the increase of partner density. For example, the following model is adapted from a previous work (81) which falls into this category. In addition, if no decreasing density is captured, this model will stabilize at its coexistence state with any positive initial density for both populations.

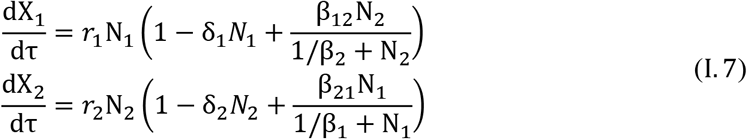

Due to the limitations of previous mutualism models in studying the transition between coexistence and collapse, a more systematic way of modeling mutualism is required to both ensure the basic logic of mutualism and capture the transition between stable coexistence and collapse.

## II Building various mutualism models

The diversity of mutualistic systems impedes the formulation of a general mutualism model and the general proof of a single criterion. We generated various mutualism models to reflect the diversity of mutualistic systems in nature and examine how diverse the coexistence criteria are. We also aim to investigate whether there exists an invariant form that is preserved regardless of specific model implementations. If such an invariant form exists, it will then reveal a fundamental coexistence criterion that is originated from the basic logic of mutualism.

### A. Incorporation of stress

We explicitly define stress as a reduction of growth rate or productivity of biological systems, which is consistent with previous works (85–88). Stress is universal in biology because it is present whenever growth rate is lower than the optimal growth rate. In mutualistic systems, many studies have shown that stress factors alter the basal fitness of individual mutualists. These factors include nutrient limitation (89), rising temperature (90, 91), rising CO_2_ levels (92), invasive species (93), etc. Beyond benefit and cost, it is known that these stress factors also determine mutualistic outcomes (94).

To study quantitatively how stress affects mutualistic outcomes, we include stress as a model parameter that reduces the growth rate or carrying capacity of individual mutualists in various biotic/abiotic contexts. In previous models, stress has been included as linear turn-over rate (66, 89, 95–97) or intraspecific competition (69, 70). Although these studies have more specific terms describing the downward pressure they capture, we use “stress” as an appropriate umbrella term.

Stress, thus defined, plays an essential qualitative role in mutualism. First, an absence of stress would mean that populations operate at maximum fitness. Such populations could not benefit from mutualism, because they would already be operating at their optimal level. Second, in mutualistic models the downward pressure can resolve the unrealistic exponential growth of mutualistic partners by imposing an upper limit to mutual benefit (78). Third, using stress can dissect the baseline fitness of individual mutualists caused by biotic/abiotic factors from the effect of mutualistic interaction on fitness.

### B. Model assumptions

The key step of generating various mutualism models is to establish the set of assumptions the models should follow. We first start off with the most apparent aspects of mutualism: benefit and cost. These two aspects lead to two assumptions:

e) Benefit shall *increase* growth rate or carrying capacity and is positively dependent on partner density.
f) Cost shall *decrease* growth rate or carrying capacity.

To study the transition between coexistence and collapse in mutualism systems, the models must be able to simulate collapse, leading to the third assumption where we explicitly introduce stress to achieve negative growth. In addition to generating negative growth, stress has a physical meaning which is the difference between maximal fitness and baseline fitness in the absence of the partner:

g) Stress shall produce negative growth of populations with some parameter combinations.

The fourth assumption follows assumption c) to reinforce the effects of benefit and cost in mutualistic interaction:

h) Negative growth of a population shall be potentially counteracted by benefit provided by a partner, but is further strengthened by cost.

Even when all the above assumptions are satisfied, we still need to verify that the models do not generate unbounded growth with *any* parameter set.

### C. Modifications of the logistic growth equation

After establishing a minimal set of assumptions, we then need to establish a systematic way of generating diverse models.

Mathematically, there are infinite possible implementations of a mutualistic system. We attempt to cover the different *common and plausible* forms of kinetic models that have been adopted in previous studies. Previous models have captured benefit, cost and stress in many ways. The following is a short summary of how benefit, stress and cost are modeled previously that serve as building blocks for our own models. Many models use Hill equations to capture saturating benefit (66, 81, 89, 98, 99). Cost is also an aspect that is widely modeled, which can be implemented many ways (81, 100), such as independent or dependent of partner density. Although stress is not often generally discussed, one possible form of stress is selfregulation, also called density dependence or inter-population competition, which often appears in large-scale models (69–71). Linear death rate (often imposed by dilution) is also another common form of stress (66, 89, 95–97).

Inspired by these observations, we first examined different methods to modify a logistic growth equation. Consider a basic logistic growth equation:

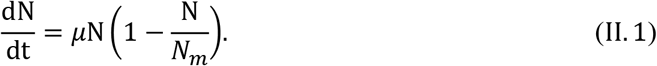

This equation only has two parameters, growth rate *μ* and carrying capacity *N_m_*. If we can derive a non-dimensionalized logistic growth equation, it can be rewritten as:

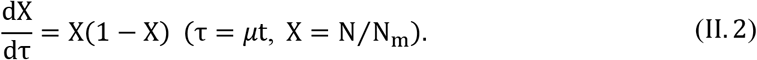

The followings are common modifications of the above equation:

1. The growth rate can be modified by multiplying the right-hand-side with a constant:

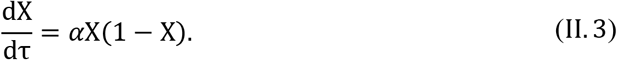 This equation modifies the growth rate from 1 to *α*, where increasing *α* increases growth rate and leaves the carrying capacity the same. *α* can take any real number in this case.
2. The carrying capacity can also be modulated, leaving the growth rate unchanged:

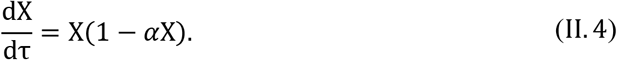 The carrying capacity becomes 1/*α* and *α* > 0. This formulation does not generate negative growth.
3. The “1” in the logistic growth equation can also be modified:

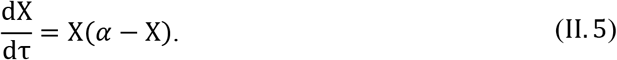 It can be rewritten as the same form of logistic growth equation:

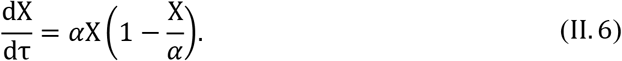 In this case, both growth rate and carrying capacity are scaled by a factor of *α*. *α* can take any real number because if the carrying capacity is negative, the growth rate will also be negative and thus the model will only generate bounded growth.
4. A first order term can be added to the equation:

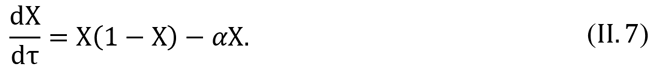 Although this modification is equivalent to the previous one, it is worth noting since this form is commonly used to represent death rate or turnover rate.

The above analysis demonstrates that 1, 3, and 4 are robust modifications that provide consistent model modulations with any real value of *α* parameter. Following this analysis, we will then model stress, benefit and cost at these three locations in the logistic growth equation to capture the assumptions presented above for mutualistic interactions.

### D. Specific implementations of benefit, stress and cost

To capture the model assumptions of a), b) and c), we used the following formulations to modify simple logistic growth equations at location 1, 3 or 4 mentioned above to capture the logic of mutualistic interaction.

#### a) Benefit shall be positively dependent on partner density

This assumption can be modeled at all three locations by

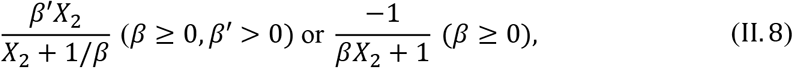

where *X_2_* represents partner density (same as below) and *β* and *β’* are measures of strength of benefit. In both cases, the benefit function is bounded, which facilitates the stabilization of population densities (73).

#### b) Cost shall decrease growth rate or carrying capacity

We use *ε* as a measure of level of cost. Specifically, this is implemented at location 1 or 3 by multiplication

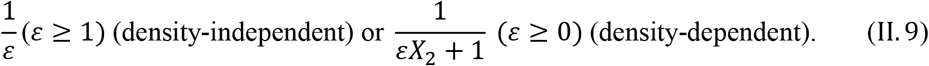

Cost can also be implemented as a turnover rate at location 4

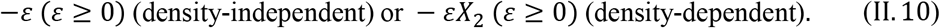

#### c) Stress shall produce negative growth of populations with some parameter combinations

We use *δ* as a measure of stress. Specifically, this is implemented by modifying the logistic growth equation at location 1 or 3 by

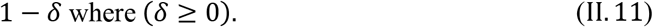

Stress can also serve as a turnover rate at location 4

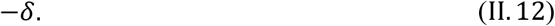

Two or more of the parameters can share one location, so we permutated the three factors in the three locations and systematically generated 81 models (Table S1). The 81 models are then checked against assumption d) and verify that the models do not generate unbounded growth. See Supplementary Materials for the MATLAB code that investigates the model assumption d) and unbounded growth behavior. Out of the 81 models, 48 satisfy all 4 assumptions and do not generate unbounded growth with positive parameter values and positive initial densities.

## III Derivation of diverse criteria

This section demonstrates the diversity of criteria with various model formulations. We first use an example model to serve as the base model to demonstrate a detailed process to derive coexistence criteria. We then show the process of building various mutualism models with more complexity and deriving or approximating the corresponding criterion.

### A. An example

Consider the following model as our base model for the analytical analyses:

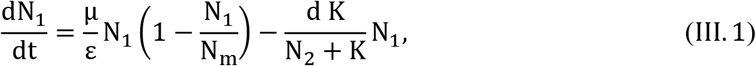

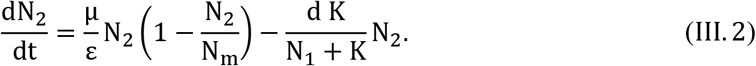

The non-dimensionalized version of the model is (model 21 in Table S1):

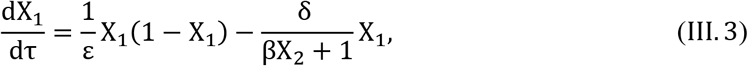

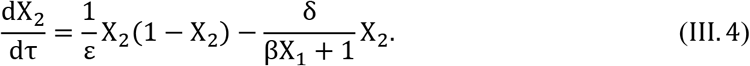

Where τ = *μ*t, X_1_ = 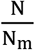, δ = 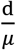, β = 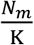. The ranges of the parameters are: *ε*≥ 1, δ≥ 0 This system has 5 fixed points:

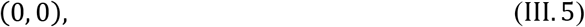

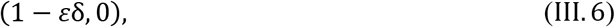

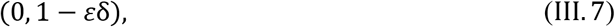

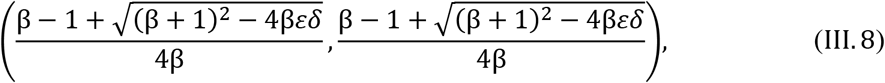

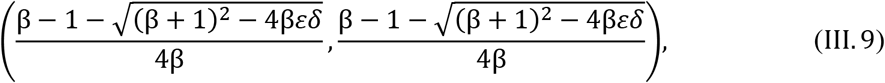

By calculating the Jacobian, we found that the 4^th^ fixed point (III.8) represents stable coexistence. The 5^th^ fixed point is a saddle point and is unstable. For the 4^th^ fixed point to be in the first quadrant of the Real domain, 2 conditions must be satisfied at the same time:

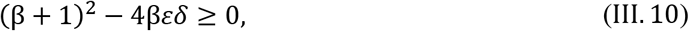

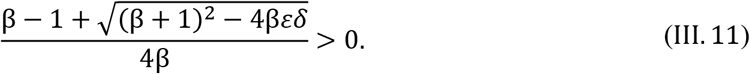

The first condition (III.10) derives the coexistence criterion 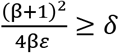 and when this criterion is satisfied, the second condition is automatically satisfied if β > 1. When β ≤ 1, only facultative mutualism can be captured. Bifurcation analysis of the system shows Allee effect and how final density increases with increasing β, decreasing *δ* or decreasing *ε*.

### B. Summary of criteria derived from symmetric models

Symmetric models in general generate interpretable analytical solutions. In Table S1, all the models are assumed to be symmetric between two populations. We found that these symmetric models already generate a diverse set of coexistence criteria.

Most of these criteria are derived using the same logic used for the base model presented above, except for the criteria of model 1, 4, 28, 31, 55 and 58 in Table S1, which are derived from setting the analytical solution of the saddle point lower than the initial density X_0_. This is because the steady states that represent coexistence for these models are constants, whereas the saddle points are modulated by model parameters. For a detailed example of the derivation, refer to section III.C.2.

In natural mutualism systems, cost can also scale with its partner’s density (100). In general, we observed that inclusion of density-dependent cost increases the complexity of the criteria. In addition, mutualists can also compete in within the same niche (101). In this case, the effective benefit term in general decreases from the model without competition, indicating the criteria also lumps the effect of competition.

### C. Criteria from models with additional complexities

To relax some additional assumptions, such as complete symmetry, we also included additional levels of complexities to the base model by including asymmetry and turnover rate:

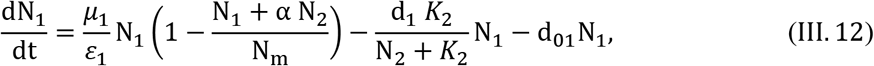

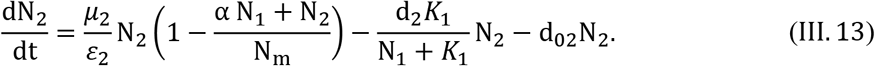

After non-dimensionalization, the model becomes:

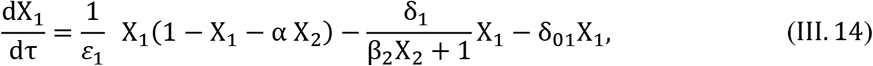

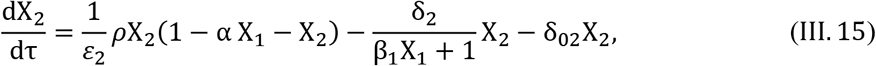

where τ = μ_1_t, X_1_ = 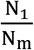,ρ = 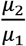,δ_1_ = 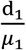,β_1_ = 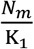, X_2_ = 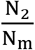,δ_2_=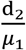,β_2_ = 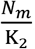.

#### 1. Criterion with two populations competing for resources

Our base model assumes the two populations having separate carrying capacities. However, in natural settings, mutualists often share resources (83). We can impose competition between them by adding *–αX_2_* or *–αX_1_* to location 3 in logistic growth equations of X_1_ and X_2_, respectively (number 82 in Table S2):

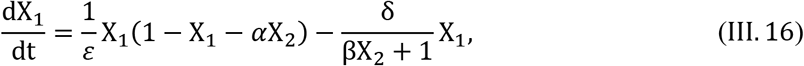

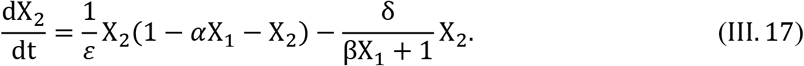

Using the same method described in section III.A, we derived the criterion of this model: 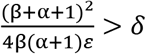

#### 2. Criterion considering initial density

For this analysis, we used a symmetric model (number 83 in Table S2):

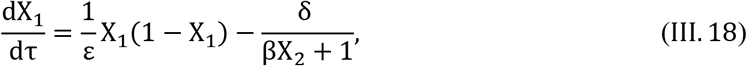

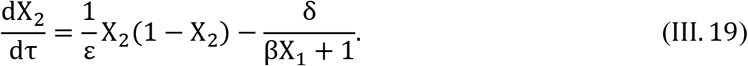

Because most mutualism models generate Allee effect, coexistence can also be affected by initial density (*x*_0_). However, we can derive a benefit term that depends on the initial density *x*_0_(for both X_1_ and X_2_) to predict deterministically whether the two populations coexist or not. To coexist, *x*_0_ needs to be greater than the saddle point:

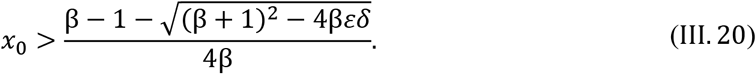

If we rearrange this inequality, we get

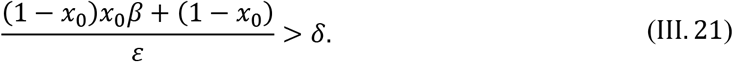

#### 3. Criterion with asymmetric growth rate, cost, stress, and benefit

Asymmetric parameters of the mutualism are more realistic in capturing real-world mutualism systems, so we relaxed the assumptions that cost (*ε*), rescue strength (β) and stress (δ) are symmetric for both populations. The asymmetry of growth rate is captured by *ρ*. In the following model (number 84 in Table S2), we assumed X_1_ and X_2_ share the same carrying capacity. We found that separated carrying capacity also yields similar results.

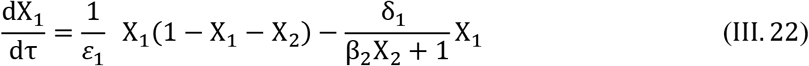

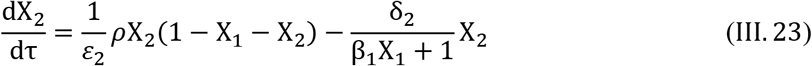

This model has 5 fixed points:

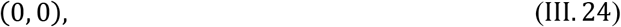

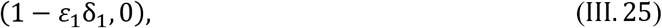

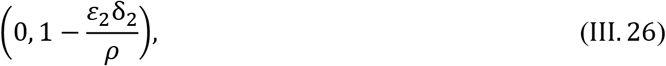

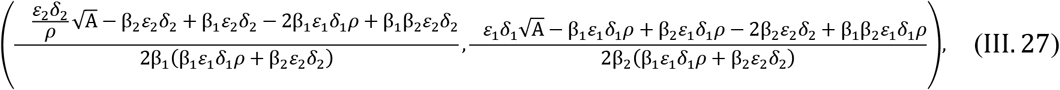

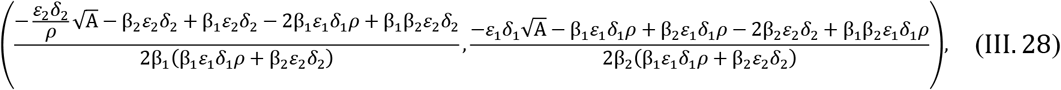

where A = ρ^2^(β_1_β_2_ + β_1_ + β_2_)^2^ — 4β_1_β_2_ρ(β_1_ε_1_δ_1_ρ + β_2_ε_2_δ_2_).

Use the same logic presented in section III.A, the following criterion is a necessary condition for the 4^th^ fixed point to be present in the positive Real domain:

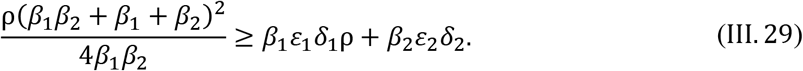

This criterion can also be rewritten as 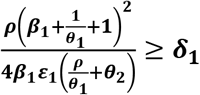 where the asymmetry is captured by 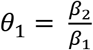 and 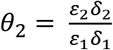. Note that we specifically used X_1_ as the reference population.

#### 4. Criterion with turnover rate and asymmetric growth rate

If we consider both the asymmetry in growth rate (*ρ*) and a turnover rate *δ*_0_, the model becomes (number 85 in Table S2):

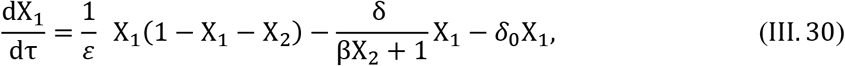

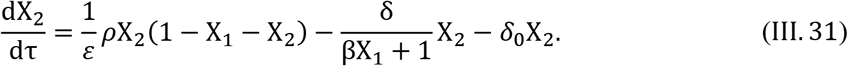

The analytical solution of this model is complex and involves solving 4^th^ order polynomials. However, we can introduce the concept of correction terms to approximate the criterion. When we only add *ρ* in the model and assume *δ*_0_ = 0, we get the following criterion:

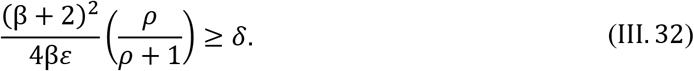

In addition, when we only add the *δ*_0_ term in the model and assume *ρ* = 1, we get

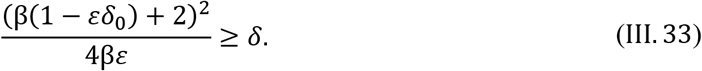

We hypothesized that the criterion with both correction terms (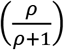) and (1 – *εδ*_0_) can approximate the criterion for model III.30-31. The criterion for *δ*_0_ > 0 is:

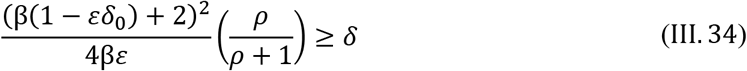

The accuracy of this criterion is evaluated by substitution of variables:

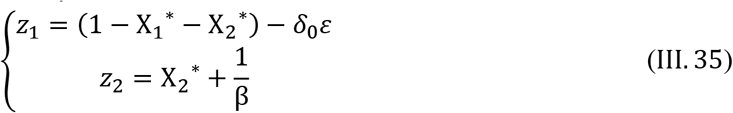

where X_1_* and X_2_* represent fixed points of the model. 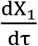 = 0 and 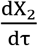 = 0 can then be written as

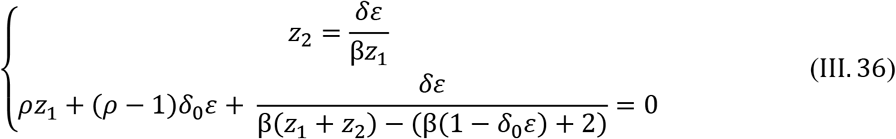

After substitution of *z_2_* with *z_1_* in the second equation, we get

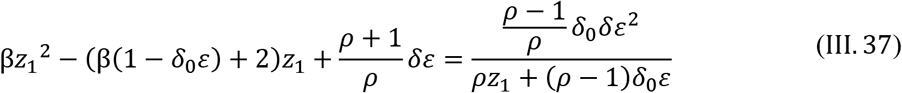

The left-hand side is a quadratic equation of has *z_1_*, which has Δ= (β(1 – *δ_0_ε*) + 2)^2^ – 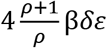.

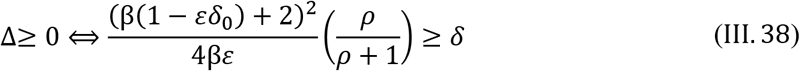

Thus, we know that the criterion is more accurate when 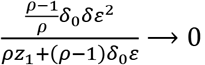. The approximated criterion is accurate when *ρ* = 1.When *ρ* → +∞ and *δ*_0_ → 0, the criterion will also provide good approximations.

#### 5. The criterion for N-mutualist systems

The following model is used to determine the criterion with N-mutualist model:

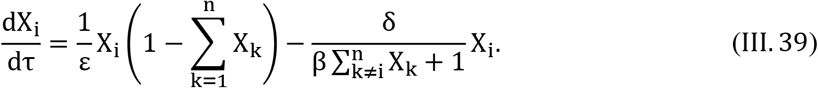

The non-trivial steady state can be solved by solving the following equation for X^*^:

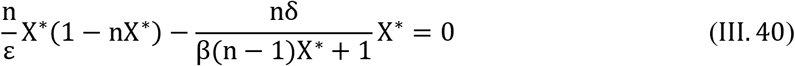

Using the same strategy in III.A, we derive the criterion for coexistence:

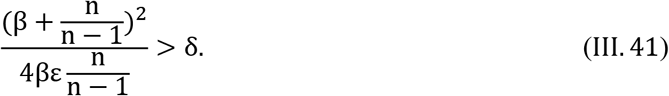

where benefit becomes a function of both β and n. This result suggests that mutualism system can tolerate higher stress levels with increasing number of mutualists.

#### 6. Mutualism model including cheater exploitation

The mutualism model including cheaters is adapted from III.22-23:

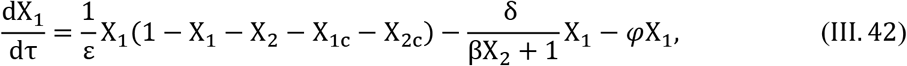

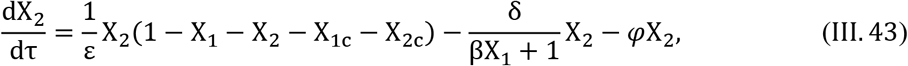

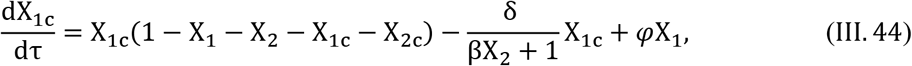

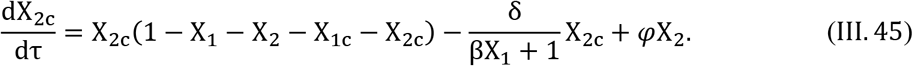

We assume 1) cheaters and cooperators share the same carrying capacity, 2) cheaters accept benefit produced by cooperators but do not provide benefit, thus do not experience cost, and 3) there is a constant transfer rate *(φ)* from cooperator to cheater, representing mutation from the cooperator phenotype to cheater phenotype. Although this model does not generate stable coexistence of the cooperators, the time it takes for cheaters to take over the cooperators can serve as a metric for how stable the mutualistic system is.

#### 7. Model structures that generate lower boundary for stress

We notice that to coexist, some model formulations not only require an upper boundary of *δ*, but also require a lower boundary as well. If *δ* is lower than a threshold, the system can be dominated by the population that is more fit. However, this lower boundary occurs due to the system dynamic shifting from a mutualism-dominated mode to a competition-dominated mode.

There are two types of system dynamics that can lead to a loss of partner. One is due to high stress where the weaker partner will go extinct and the stronger partner will suffer from the loss of its partner. The other is due to lack of stress where the system shifts to a competition-dominated interaction where the fitter partner survives better by excluding its partner. In our study, we only focus on the first type of partner loss since the second type is a dynamic of competitive systems instead of mutualistic systems.

In general, we observed that a model needs to be both asymmetric and potentially competitive to create a scenario where the fitter population excludes the weaker population. The lower boundary for *δ* decreases when the weaker population gives out more benefit to its partner or reduction of competition. This observation can potentially explain why some mutualistic systems collapse when external stress is reduced (89, 102).

## IV. Theoretical generality of the simple metric

To establish the generality of the theoretical criterion, we verified that it is applicable to both symmetric and asymmetric mutualistic systems. We also want to test that the predictive power of the predictor is maintained when the partners are obligatory or facultative. Because mutualists can compete for the same resources in nature, we also verified that coexistence of partners that compete for resources also can be depicted by the criterion. We show some typical results of the predictive power of the criterion in these cases with Fig. S2.

### A. Establish the general predictive power of *B/δ*

Fig. 1f is generated using the model presented by equation III.16-17, where ε ∈ [1,1.2], δ ∈ [0, 2], β ∈ [2, 5] and *a* = 1. The positive trend holds even when the mutualism is facultative (δ < 1).

Fig. S2a is also generated using the model presented by equation III.16-17. Both are generated with β ∈ [2, 10], δ ∈ [1.2,2], ε ∈ [1,1.2] and *a* = 1. The panel on the left assumes X_1_ and X_2_ have the same initial density which is a uniformly distributed between 0 and 0.5. The panel on the right assumes changing of individual initial densities in a linear fashion while maintaining the sum of initial density of X_1_ and X_2_ at 1.

Fig. S2b is generated with the model presented above in equation III.42-III.45, where *φ* = 10^−3^, β ∈ [5, 20] δ ∈ [1.2,1.3], and ε ∈ [1.2,1.3]. The initial densities are X_10_ = 0.1, X_10_ = 0.1, X_1c_ = 0 and X_2c_ = 0. The time to cheater exploitation is quantified by the first time point where the total density of cooperators drops below their initial total density due to overwhelming competitions from the cheater populations. Other time to cheater exploitation in this study is quantified using the same method.

Fig. S2c is generated by

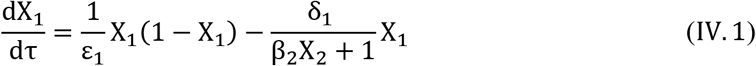

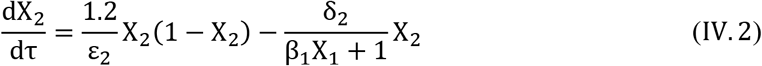

Where the left panel uses parameter values of β_1_ = 2, β_2_ ∈ [2,5], δ_1_ = 1.2, δ_2_ ∈ [0.5,2], ε_1_ = 1.1, and ε_2_ ∈ [1,1.2]. The initial densities are X_10_ = 1 and X_20_ = 1. The right panel uses parameter values of β_1_ = 2, β_2_ ∈ [2,5], δ_1_ = 0.8, δ_2_ ∈ [1,2], ε_1_ = 1.1, and ε_2_ ∈ [1,1.2]. The initial densities are X_10_ = 1 and X_20_ = 1. This panel specifically tests the cases where the survival of the partners is fully dependent on each other.

Fig. S2d is generated by III.22-III.23, where the left panel uses parameter values of *ρ* = 1.2, β_1_ = 2, β_2_ ∈ [5,10], δ_1_ = 1.2, δ_2_ ∈ [0.5,2], ε_1_ = 1.1, and ε_2_ ∈ [1,1.2]. The initial densities are X_10_ = 1 and X_20_ = 1. The right panel uses parameter values of β_1_ = 2, β_2_ ∈ [5,10], δ_1_ = 0.8, δ_2_ ∈ [1, 2], ε_1_ = 1.1, and ε_2_ ∈ [1,1.2]. The initial densities are X_10_ = 1 and X_20_ = 1. This panel specifically tests the cases where one mutualist’s survival is not fully dependent on the other.

Fig. S2e is generated using the same model as the above. Where the parameter values are β_1_ = 5, β_2_ ∈ [2,10], δ_1_ = 1.2, δ_2_ ∈ [1.2,2], ε_1_ = 1.1, and ε_2_ ∈ [1,1.2]. The sum of initial densities is kept at 0.3 and the values of X_10_ and X_20_ are changed in a linear fashion. 1000 different combinations of parameters are used and 50 different X_10_: X_20_ simulations are performed with each parameter combination to calculate the probability of coexistence.

Fig. S2f is generated using the following model, which is based on the symmetric model presented in equation III.42-45, while adding asymmetry:

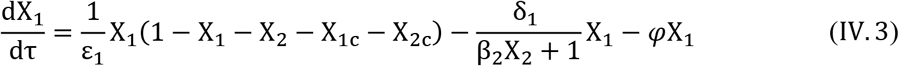

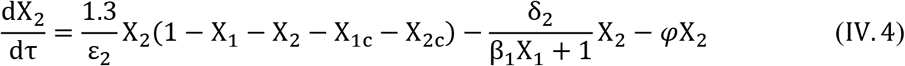

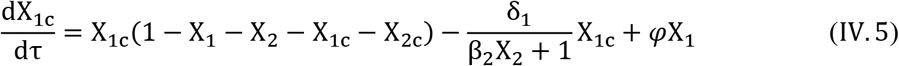

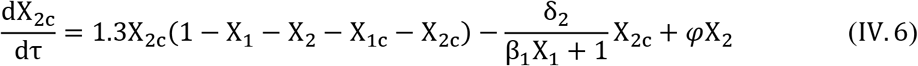

Where *φ* = 10^−3^, β_1_ = 4, β_2_ ∈ [40,60], δ_1_ = 1.8, δ_2_ ∈ [1.4,1.5], ε_1_ = 1.4, ε_2_ ∈ [1,1.2]. The initial densities are X_10_ = 0.05, X_10_ = 0.05, X_1c_ = 0 and X_2c_ = 0.

### B. The predictive accuracy is maintained in the presence of noise

In addition to investigating the probability of coexistence when initial density is randomly distributed (Fig. S2a, e), we also explicitly modeled noise to test the effect of noise on the prediction accuracy of the criterion. We added Gaussian noise to our base model *(η*):

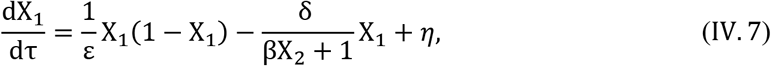

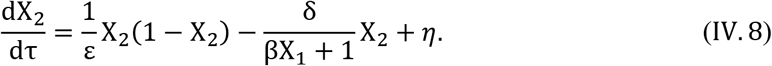

In Fig. S2g, we used the model above and the criterion tested is III.19. Multiple sets of initial densities are tested. Although we observed a decreasing accuracy with increasing noise, the prediction accuracy is robustly maintained (above 90%) even when the standard deviation of noise reaches 50% of signal strength. The high maintenance of prediction accuracy is due to the stability of the mutualistic model. Because the two stable steady states exist on opposite sides of the separatrix, only when noise pushes densities across the separatrix (against the vector field), the outcome of the system will be altered. Otherwise, noise will not influence the final steady state.

## V. Calibration using SVM

We chose SVM because it is a well-established algorithm and it only requires information of the support vectors which are usually data points along the boundary to obtain a classification boundary. Note that other algorithms can also be implemented for the same purpose. For example, when predicting probability of coexistence, logistic regression can be more suitable to directly predict probability of coexistence. See the Supplementary Materials for our MATLAB program which performs the calibration procedure using SVM.

### A. Kernels

We used four types of kernels to cover different general shapes of the *B* landscape:

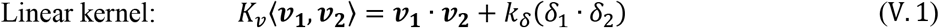

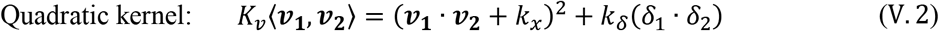

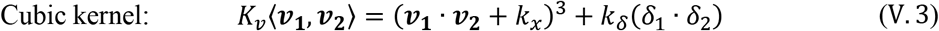

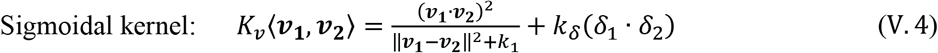

Where in equation V.4,**‖*****v_1_ – v_2_*****‖**^2^ = Σ_i_(*v*_1*i*_ – *v*_2*i*_)^2^.

These four kernels follow the same structure:

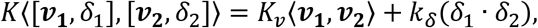

Our procedure allows any kernel structure that is supplied by the user, so other customized kernels can also be used for the calibration.

### B. Standardizing input data

Standardizing input data is essential for robust learning of *B*(***v***). Before training the model with SVM, we first standardize the system variables *v* and stress *δ* to mean 0 and variance 1.

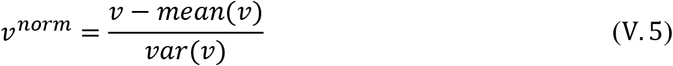

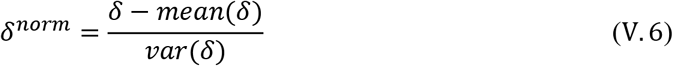

### C. From SVM output to calibrated *B(v)*

SVM will output a predictor that separates coexistence and collapse indicated in the ***Y*** vector. Our program requires that the two classes are represented by 1 and −1 in the ***Y*** vector. Using kernels V.1-4, we can write the predictor in the general form:

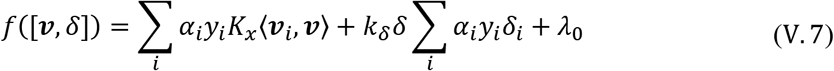

where *k_δ_* is a kernel parameter; *α_i_* represents coefficients associated with each input observation that are optimized by the SVM algorithm; *λ*_0_ is the bias that is optimized by the algorithm. If we impose *B* = *δ* at *f* ([**v**, δ]) = 0, we can obtain a function of *B* in terms of input data:

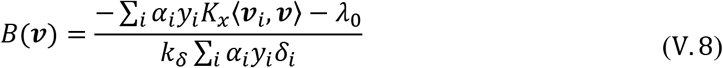

Using this function, we can obtain a value of *B* with any system variable dictated by ***v*** based on the limited number of observations used as inputs.

*B*(***v***) can sometimes have wrong directionality, meaning that *β/δ* > 1 is associated with collapse and *B/δ* < 1 is associated with coexistence. We identify these cases by calculating

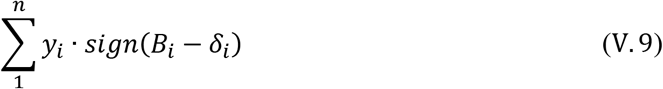

If the above expression is negative (when the trained boundary have a learning accuracy greater than 50%), the calibrated *B*(***v***) has a wrong directionality. Flipping of the *B* landscape is then required, and it is done by assuming:

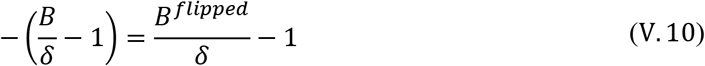

Thus,

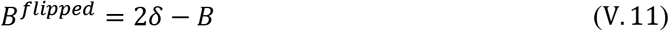

We name the output *B* landscape with properly adjusted directionality *B’*. This landscape *B’* describes the shape of the final calibrated *B*(***v***) but still needs to be rescaled for prediction purpose. The final *B*(***v***) is calculated by rescaling it back according to the mean and variance of the *δ* measurements.

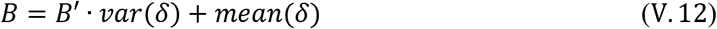

### D. Cross validation and bootstrapping

All the cross validations in this work are 10-fold cross validations. The cross-validation accuracy is represented by the average values. Bootstrapping is used to evaluate the degree of variation of quantified *B*. The same number of data points as the input data is randomly sampled from the input data with replacement. We performed bootstrapping for 500 times. The variance and relative standard deviation (RSD) are then calculated based on the 500 bootstrapped *B*(***v***) quantified with 500 sets of sampled training data. The mean cross validation accuracy, the bootstrapped variance and relative standard deviation are then used to evaluate the accuracy of the *B*(***v***) outputs.

### E. Twenty sets of simulations to establish and test the calibration procedure

To establish the calibration process, we first developed the method with simulated data where the true *B*(***v***) are known. This allows us to evaluate calibration results against the true function. We used the model presented in equation III.22-23 to conduct this set of analyses. Using *X_1_* as the reference population, the *B*(***v***) can be expressed as:

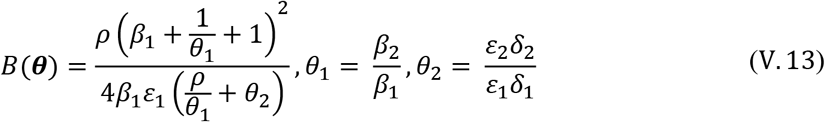

However, depending on how system variables change the model parameters *(**θ**(**v**))* in the system variable space, the true *B(**θ**(**v**))* can exhibit diverse shapes. With *v_1_, v_2_* ∈ [0,1], we constructed 20 arbitrary set of equations with different underlying functions. We also made sure the generated data have around 50:50 split of coexistence and collapse in the simulated results to reduce bias in the input data. This 50:50 split constraint also applies to the all other simulated data. Note that these functions are arbitrary and only serve the purpose of generating true *B*(***v***) that has various underlying functions. The data generated using these 20 models are included in the MATLAB code in the Supplementary Materials.

The 20 sets of ***θ**(**v**)* are grouped into 5 types with each type containing 4 different examples. The 5 types of functions that describe ***θ**(**v**)* include linear, quadratic, cubic, and Hill equation and another type of functions that are a mixture of the previous four types. As examples, the following are four sets of ***θ**(**v**)* used to generate simulation results shown in Fig. S4b.

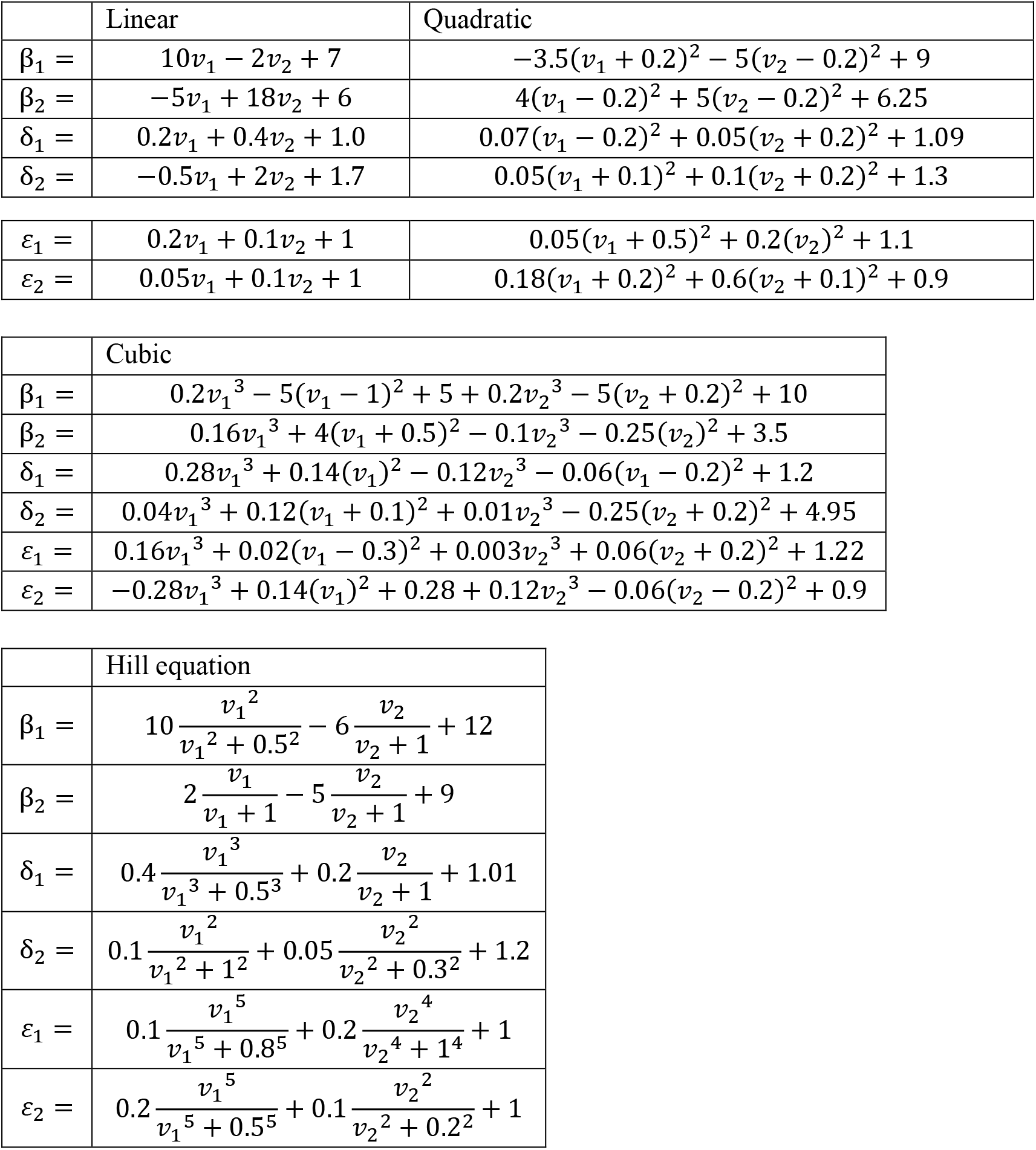

The training data are 10 × 10 on the *v_1_, v_2_* space. *R^2^* between calibrated *B* landscape *(B*_c_) and the true landscape *(B_t_)* is calculated by first performing a least square linear fit of *B_t_* with *B_c_*. This fitting process aims to get a linear transformation of *B_c_* that conforms with the scale of *B_t_* while maintaining the shape of *B_c_*. The absolute scale of the calibrated *B*(***v***) is less crucial than its shape, and an absolute scale is also challenging to obtain.

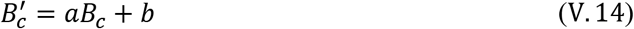

Then *R^2^* is then calculated by:

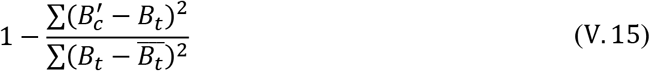

### F. Calibration with simulated data with unknown *Β*(θ): an example

We want to test whether we can apply the calibration procedure to data that are generated by an arbitrary mutualism model. The specific model structure we used to generate data in Fig. 2c is:

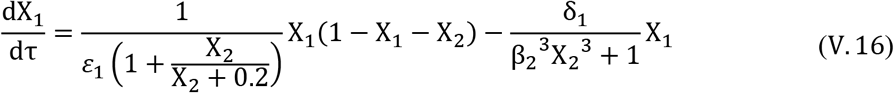

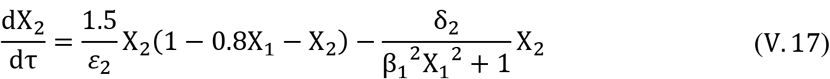

The analytical solution of this model cannot be expressed in a simple and explicit form. However, using simulated data, we can obtain an empirical function of *B* on the system variable space that allow further prediction in the system variable space. The simulations are done using initial densities of [0.01, 0.01] for 10 unit-time.

To get the probability of coexistence, we ran the model 100 times with varying ratios of initial density while keeping the total initial density constant as 0.02. The density for a population is varied in a linear fashion from 0 to 0.02 and we terminated the simulation after 10 unit-time.

To calculate how well the systems can resist cheater exploitation, we modified the equations V. 16 — 17 to account for the emergence of cheaters:

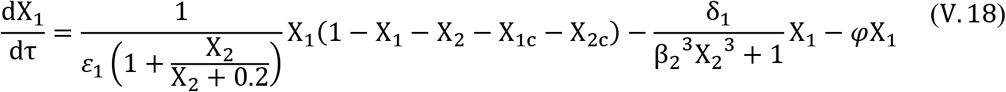

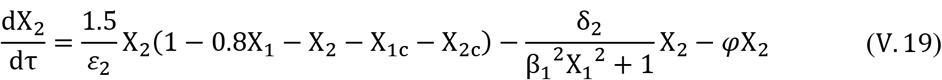

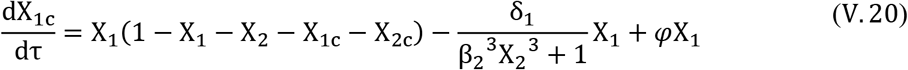

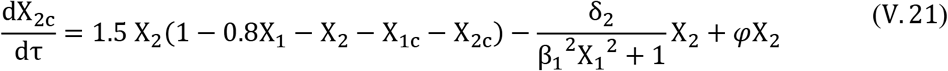

The simulations were done with *φ* = 10^−3^. The initial densities are [0.01, 0.01, 0, 0] for X_1_, X_2_, X_1c_, and X_2c_, respectively. The simulations were terminated after 1300 unit-time.

## VI. Framework generality verified by complex mutualistic systems

### A. A5-mutualist system

The data presented in Fig. 4a and Fig. S9a are generated with the following mutualism model based on the general model structure presented in equation III.39:

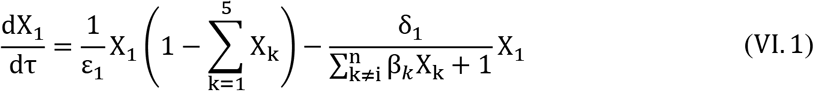

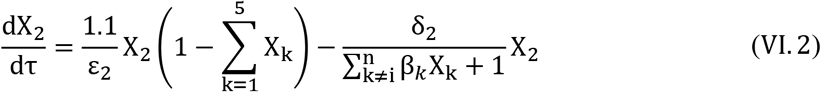

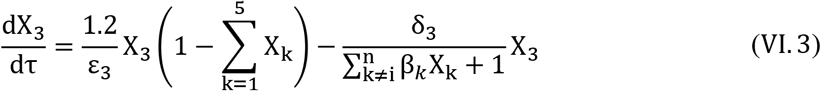

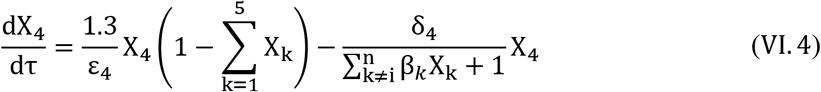

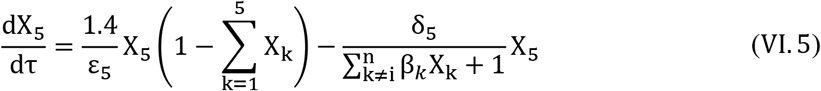

All the parameters are linear combinations of *v*_1_, *v*_2_ ∈ [0,1] and the coefficients are randomly generated. The initial densities are [0.001, 0.001, 0.001, 0.001, 0.001] for X_1_ to X_5_ respectively. The simulations were terminated after 5 unit-time.

### B. The experimental 3-member mutualistic systems

The calibration shown in Fig. 4b is done on all 384 systems with triplicates that alternatively use each strain in a system as the reference strain. The δ measurements used in this calibration is the same as the measurements presented in Fig. 3j and Fig. S8b with the pairwise systems.

### C. A mutualistic system in an oscillatory environment

The oscillatory signal is implemented as square pulses, where

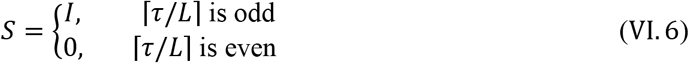

*τ* is time; *L* is the duration of the pulses and the duration between the end of one pulse and the start of the next pulse; *I* is the intensity of the signal. Throughout the simulation, δ_1_, δ_2_, β_1_, and β_2_ are affected by this oscillating signal, where

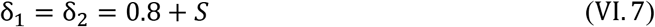

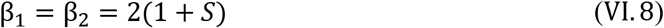

To simulate the data, we used a logarithm scale that vary L from 10^−2^ to 10^2^ and vary *I* from 10^−1^ to 10^1^. The model used to generate the set of data in Fig. 4c and the left column of Fig. S10 is presented in equation III. 22-III.23, where *ρ* = 1.5. The initial densities used are [0.2, 0.2] for X_1_ and X_2_ respectively. The simulations are performed for 500 unit-time. The theoretical *β*(*θ*(***v***)) is calculated using III.32 with δ_1_ = δ_2_ = 0.8 + *I* and β_1_ = β_2_ = 2(1 + *I*).

### D. A mutualistic system cohabiting with other populations

To generate simulated data presented in Fig. 4d and the right column of Fig. S10, we used the following arbitrary 7-population model, where X_1_ and X_2_ are mutualistic and their population dynamics are modulated by the other 5 populations.

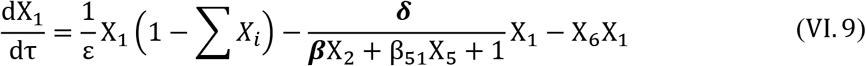

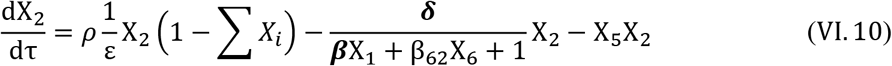

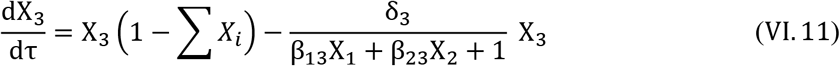

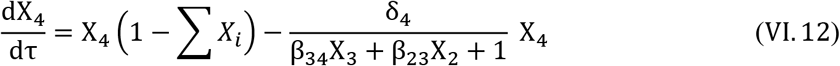

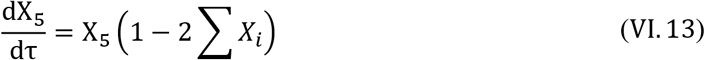

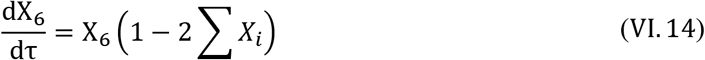

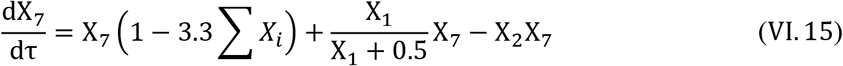

In the simulation, only *δ* and *β* are modulated by system variables *v_x_, v_2_* ∈ [0,1], while other parameters were kept constants.

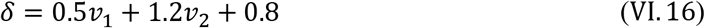

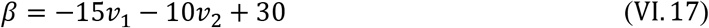

The initial densities are [1, 1, 1, 1, 1, 1, 1] for *X*_1_ to *X*_7_ respectively. The simulations were performed for 5000 unit-time. The theoretical *B(**θ**(**v**))/δ* in the right panel of Fig. S10a is directly calculated using III.32.

---

**Table S1:**
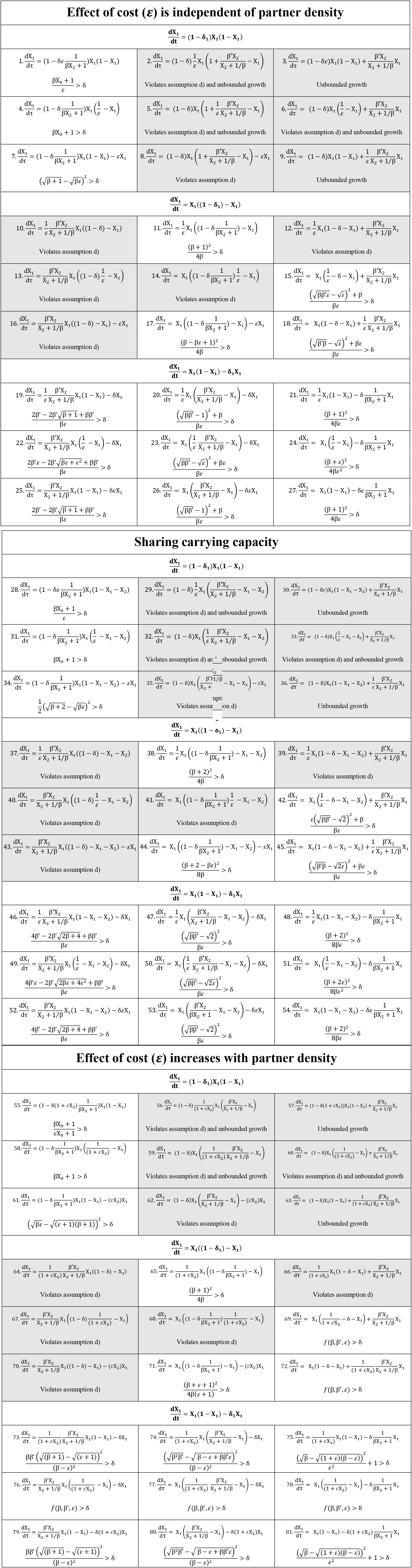
Diverse coexistence criteria derived from a set of 81 mutualism models. The formulations are based on the locations of benefit (*β, β’*), cost (*ε*), and stress (*δ*) in logistic growth equations (see section II of SI for detailed method). Each *3 × 3* block represents different locations of the *δ* term. Each column and row represent a location of the benefit and cost term, respectively. Because we assume symmetry between the two populations, only the equation representing X_1_ is shown for simplicity of presentation. Models in the first table assume cost is independent of partner density. In natural mutualism systems, mutualists can also compete for resources, so the third table contains models that describe the two partners sharing the same carrying capacity. In addition, cost can also scale with partner densities, so models in the second table add density-dependent cost as linear dependencies. Models highlighted in grey either violate model assumption (see Method section) or generate unbounded growth, or both. The rest of the 48 models satisfy the 4 model assumptions and have bounded growth. For criteria that have a long left-hand side are written as *f(β, β’, ε) > δ*. The specific forms of *f(β, β’, ε) > δ* can be found in the supplemented MATLAB code. The MATLAB code also includes all 81 models in this table and the process of testing assumptions, calculating and verifying the above criteria.

**Table S2:** Increasing model complexity increases criterion complexity. The base model is identical to model 21 in Table S1. Number 82-86 and their corresponding criteria are obtained by relaxing assumptions in model 21. Including the 48 models in Table S1, we in total obtained coexistence criteria of 52 models. Note that number 21 and number 83 used the same model structure. The detailed models and criterion derivations are shown in section III of SI. Also see Supplementary Materials for the MATLAB code of above models and the process of calculating and verifying the corresponding criteria.

| Adding complexities | Inequality | Parameters |
| --- | --- | --- |
| Base model criterion | 21. $\frac{(\beta + 1)^2}{4\beta\varepsilon} > \delta$ | $\beta, \varepsilon$ |
| Include competition | 82. $\frac{(\beta + a + 1)^2}{4\beta\varepsilon(a + 1)} > \delta$ | $\beta, \varepsilon, a$ |
| Include initial density | 83. $\frac{1}{\varepsilon}(1 - x_0)x_0\beta + \frac{1}{\varepsilon}(1 - x_0) > \delta$ | $\beta, \varepsilon, x_0$ |
| Include asymmetry | 84. $\frac{\rho(\beta_1 + 1/\theta_1 + 1)^2}{4\beta_1\varepsilon_1(\rho/\theta_1 + \theta_2)} > \delta_1$ $\theta_1 = \frac{\beta_2}{\beta_1}, \theta_2 = \frac{\varepsilon_2\delta_2}{\varepsilon_1\delta_1}$ | $\beta, \varepsilon, \rho, \theta_1, \theta_2$ |
| Include turnover rate | 85. $\frac{(\beta(1 - \varepsilon\delta_0) + 2)^2}{4\beta\varepsilon} \left( \frac{\rho}{\rho + 1} \right) > \delta$ | $\beta, \varepsilon, \rho, \delta_0$ |
| N-mutualist system | 86. $\frac{(\beta + \frac{n}{n-1})^2}{4\beta\varepsilon \frac{n}{n-1}} > \delta$ | $\beta, \varepsilon, n$ |

**Figure S1:**
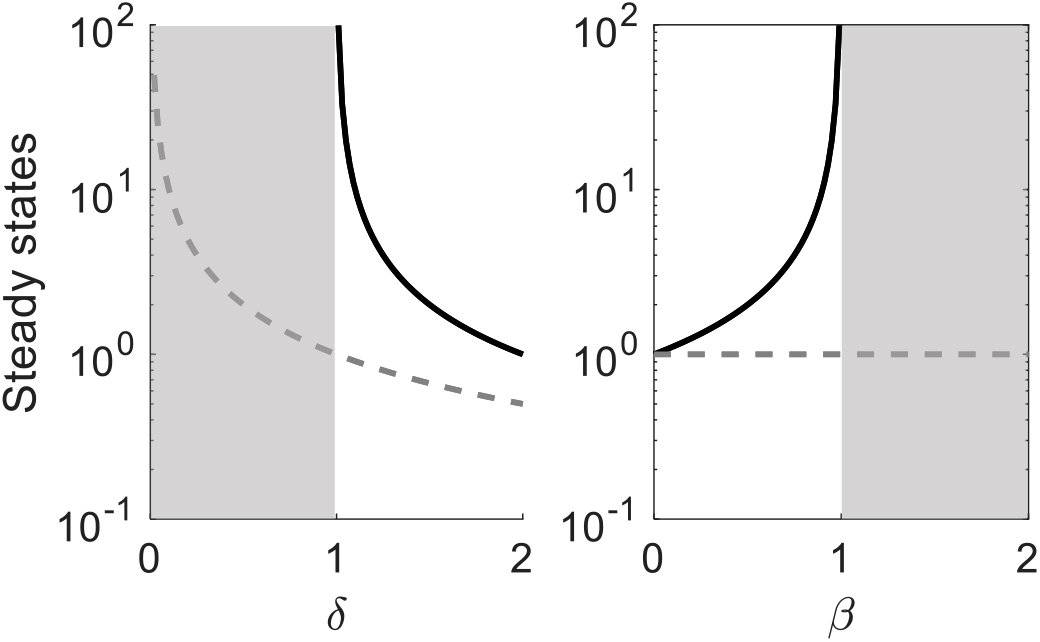
Typical bifurcation diagrams of models that create unbounded growth. Previously-developed general coexistence criteria describe the boundary between stable coexistence (white regions) and unbounded growth (grey regions). Solid lines represent stable steady state for coexistence. Grey dashed lines represent steady states of a mutualist in the absence of its partner. This diagram is generated using model:

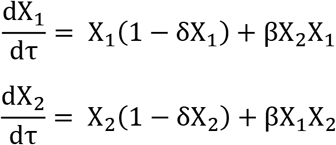

The left panel is generated with β = 1 and the right panel is generated with δ = 1.

**Figure S2:**
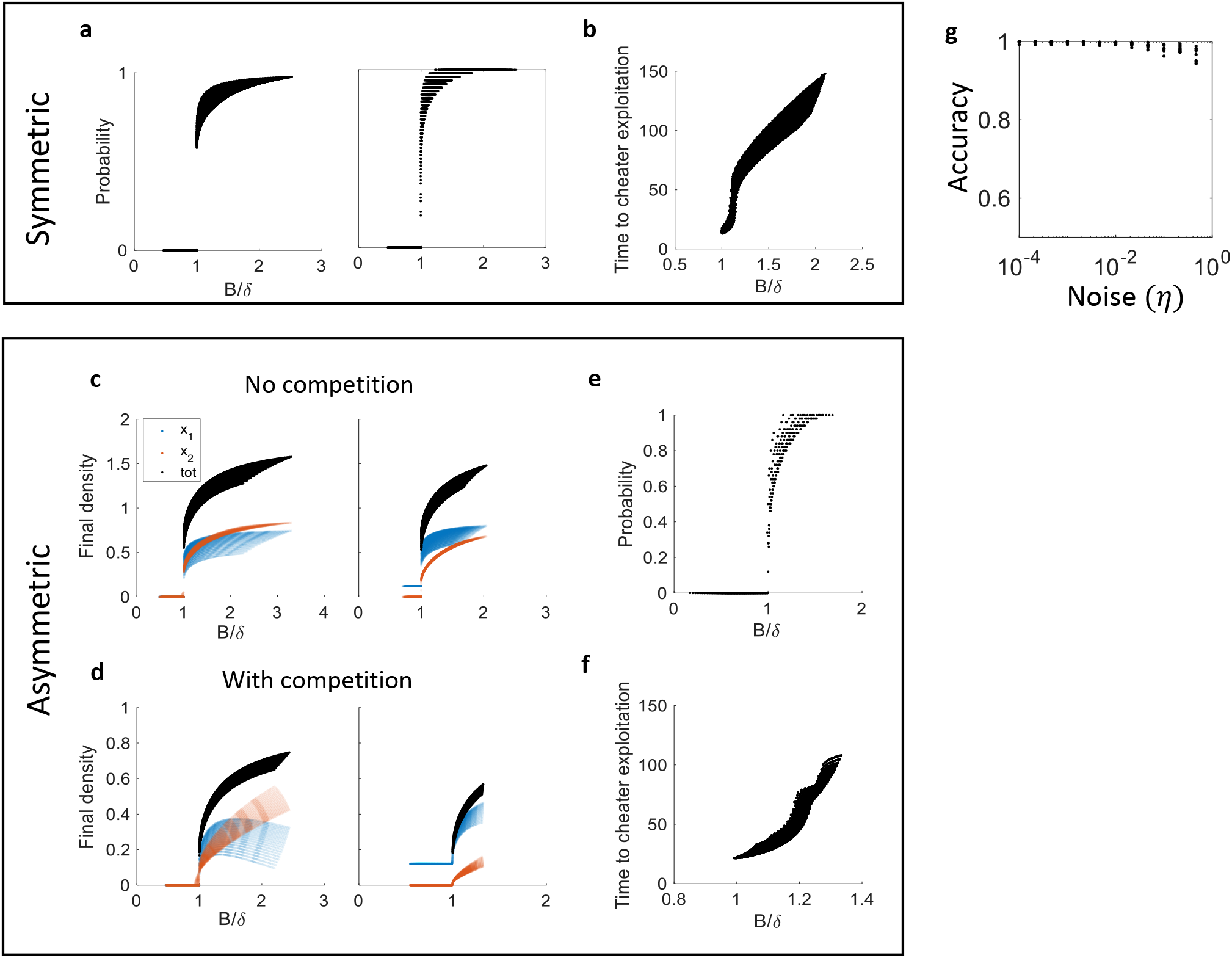
The predictor *B/δ* is a general metric for predicting performance of symmetric (panel a, b) and asymmetric mutualistic systems (panel c-f). All the traces are comprised of individual dots. Each dot represents result from one set of model parameter. **a**. Mutualistic systems have initial-density-dependent coexistence. In the left panel, we assume the same initial density of both populations that is a uniform random variable. In the right panel, we varied the ratio of the two initial densities and kept the total initial density constant. In both cases, *B/δ* is predictive of coexistence probability. **b**. When cheaters can arise in a system, *B/δ* is predictive of the time duration the system can persist before cheater exploitation. **c**. An asymmetric mutualistic system can be either obligatory (left panel), where both populations extinct when *B/δ < 1* or facultative (right panel), where one population can persist even when *B/δ < 1*. The black dots represent total density and the two colors represent the total density of the two partners. The same representations are used in panel d. **d**. The predictive power of *B/δ* for total density also holds for asymmetric systems that share carrying capacity. **e**. *B/δ* is predictive of probability of coexistence for asymmetric systems. **f**. The predictive power of *B/δ* for resistance to cheater exploitation also holds for asymmetric systems. Please refer to SI for models and parameter values used in generating this figure. **g**. The accuracy of criterion is robustly maintained after addition of extrinsic Gaussian noise. X axis indicates the standard deviation of the Gaussian noise. Note that the noise is significant considering the maximum total density of the system is 1. Please see Supplementary Text section IV.B for the simulation details.

**Figure S3:**
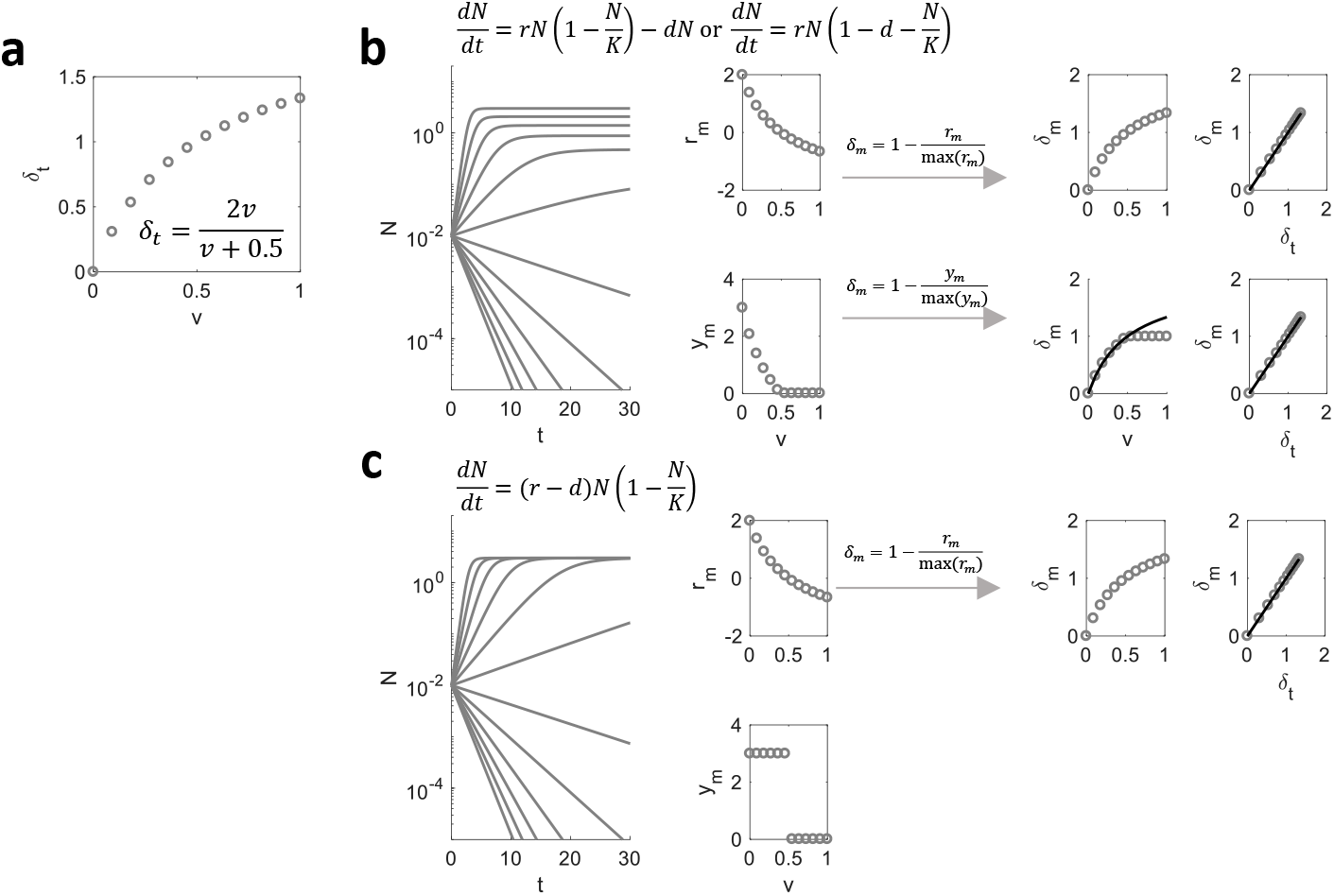
A general method for quantifying *δ*. Regardless of different model formulations, *δ* can always be measured using either growth rate or final population size of an individual population. **a**. We simulated the growth of a population that is modulated by a death term *d*. Stress *δ* is simply *d* normalized by growth rate *r*. In the simulations, the true *δ (δ_t_*) depends on an independent variable ***v***. **b**. One type of model structure has *δ* modulating both growth rate (*r*) and yield (*y*, final density). *N* is the population size and *K* is the carrying capacity. Based on the growth curve or final population size, measurements of growth rate (*r_m_*) and yield (*y_m_*) can be obtained when varying ***v***. The *δ* measurement (*δ_m_*) can be calculated by 1 — 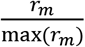 or 1 – 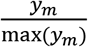. Note that since *y_m_* is always non-negative, 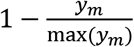 is always positive, but *δ* can be greater than 1 when the population size decreases. To address this discrepancy, extrapolation of *δ_m_* is needed to capture death rate (the black trace in the *δ_m_* versus ***v*** plot). **c**. In another type of model formulation, *δ* only modulates growth rate. Simulation shows that *y_m_* either equals to carrying capacity or 0. Thus, in this case, only *r_m_* can be used to calculate *δ*. The right-most panels show the comparison between the measured *δ (δ_m_*) and true *δ (δ_t_*). In all cases, *δ_m_* is a good approximation of *δ_t_* (the solid black line represents *δ_m_ = δ_t_*).

**Figure S4:**
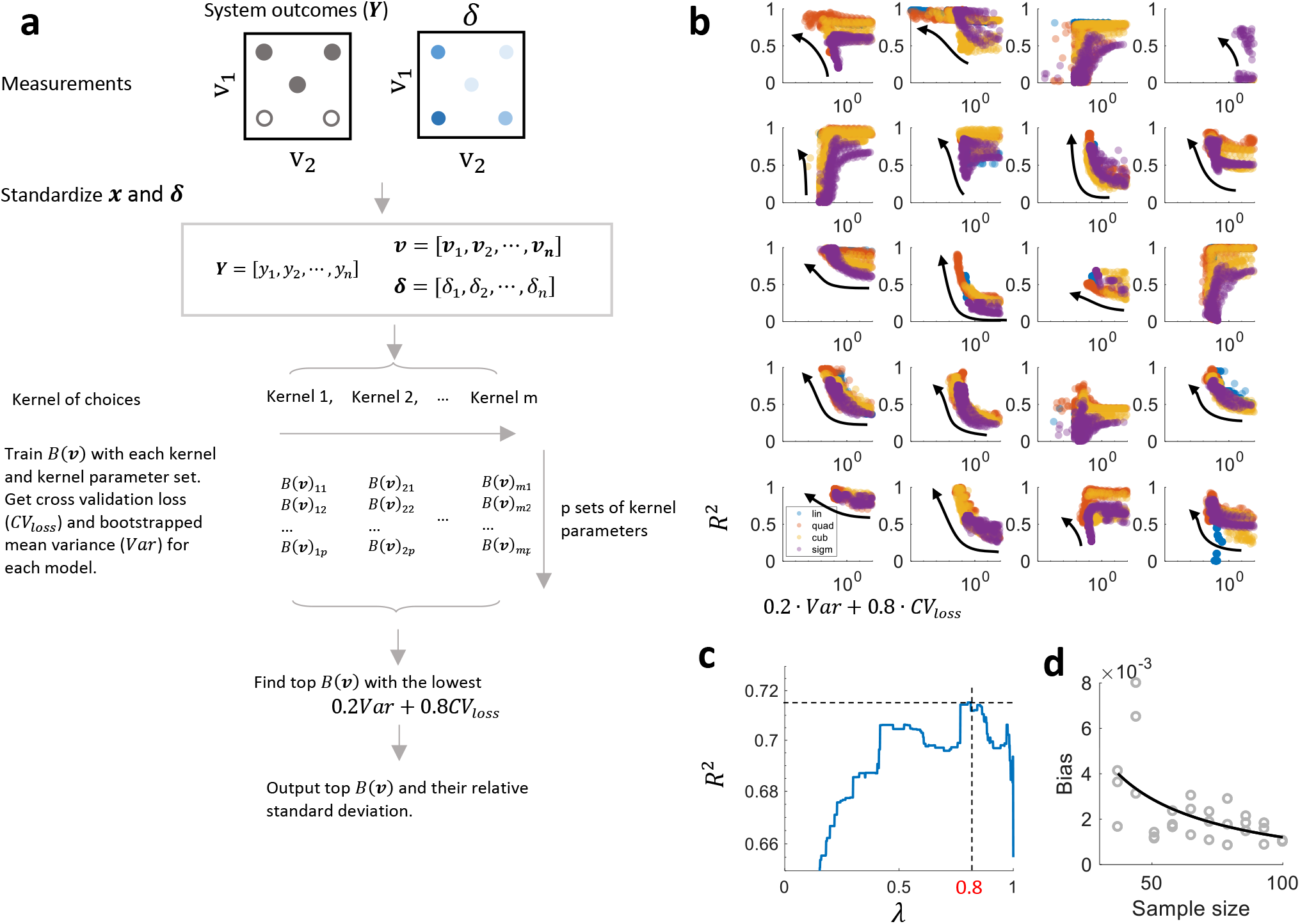
The calibration procedure using SVM. **a**. Measurements are first arranged in matrix forms and each standardized to have mean of 0 and standard deviation of 1. The standardized data are used as inputs to train SVM models with different kernels and kernel parameters. 10-fold cross validation accuracy loss (*CV_loss_*) and variance (*Var*) are calculated for each trained *B*(***v***). Top *B*(***v***) are selected based on minimizing the metric 0.2 · *Var* + 0.8 · *CV_loss_*. We then output the top *B*(***v***) (usually 5) and their corresponding relative standard deviation (*RSD* = *variance/mean*) obtained by bootstrapping to evaluate the confidence of the calibrated *B*(***v***) **b**. To show that 0.2 · *Var* + 0.8 · *CV_loss_* is a consistent indicator of high correlation (indicated by R^2^ values) between *β(v*) and the true β, we constructed 20 models that have known *B*(***v***) (each subplot represents one model). Each dot represents the result of one set of kernel parameter and the colors represent the type of kernel used. These models show that minimizing 0.2 · *Var* + 0.8 · *CV_loss_* leads to higher R^2^ values (indicated by the black arrows). See SI for details of the specific model and model parameters used to generate data in this panel. **c**. 0.8 in the metric is picked by sweeping *λ* in (1 — *λ*) · *Var* + *λ* · *CV_loss_*. Mean R^2^ calculated using the top 3 *B*(***v***) of each 20 model varies with *λ* values. The max mean R^2^ occurs at *λ* = 0.8. **d**. Bias decreases exponentially with increasing sample size. This analysis is done using the model corresponding to the third subplot in the first column of panel b. A subset of the 100 data points is randomly sampled without replacement to train *B*(***v***) and the bias of *B*(***v***) is calculated. The black trace represents the data points fitted to the function *Bias* = *a* · (*Sample size*)^−α^ + *b*.

**Figure S5:**
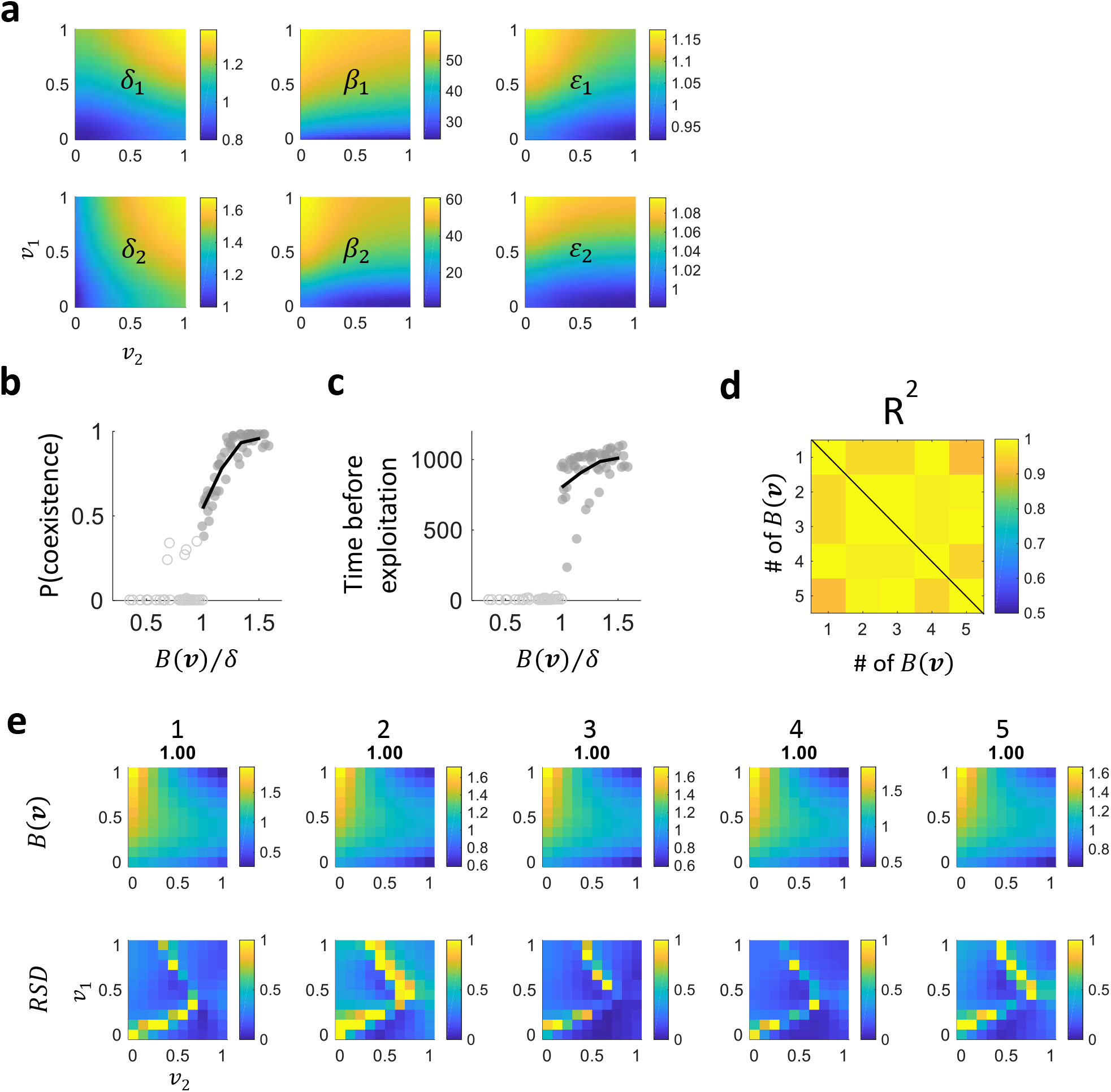
Application of the simple rule to a complex model with no explicit form of *B*. **a**. *δ*_1_, *δ*_2_, *β*_1_, *β*_2_, *ε*_1_, and *ε_2_* vary with ***v***_1_ and ***v***_2_. **b**. The calibrated *B*(***v***) along with *δ* is predictive of probability of coexistence. To get the probability of coexistence, at each (***v***_1_,***v***_2_), we ran 100 simulations with 100 different ratios of initial densities while keeping the total initial density the same. **c**. *B(**v**)/δ* is also predictive of how well the system can resist cheater exploitation. The y axis indicates the time the system can persist before the exploitation of cheater populations. **d**. The high pairwise R^2^ values of the top 5 *B*(***v***) indicate high confidence of the calibration results. **e**. The top 5 *B*(***v***) from the calibration procedure. The title for each *B(**v***) indicates its index corresponding to panel d and its corresponding training accuracy. Each *B*(***v***) can also be evaluated based on their bootstrapped relative standard deviation (*RSD*).

**Figure S6:**
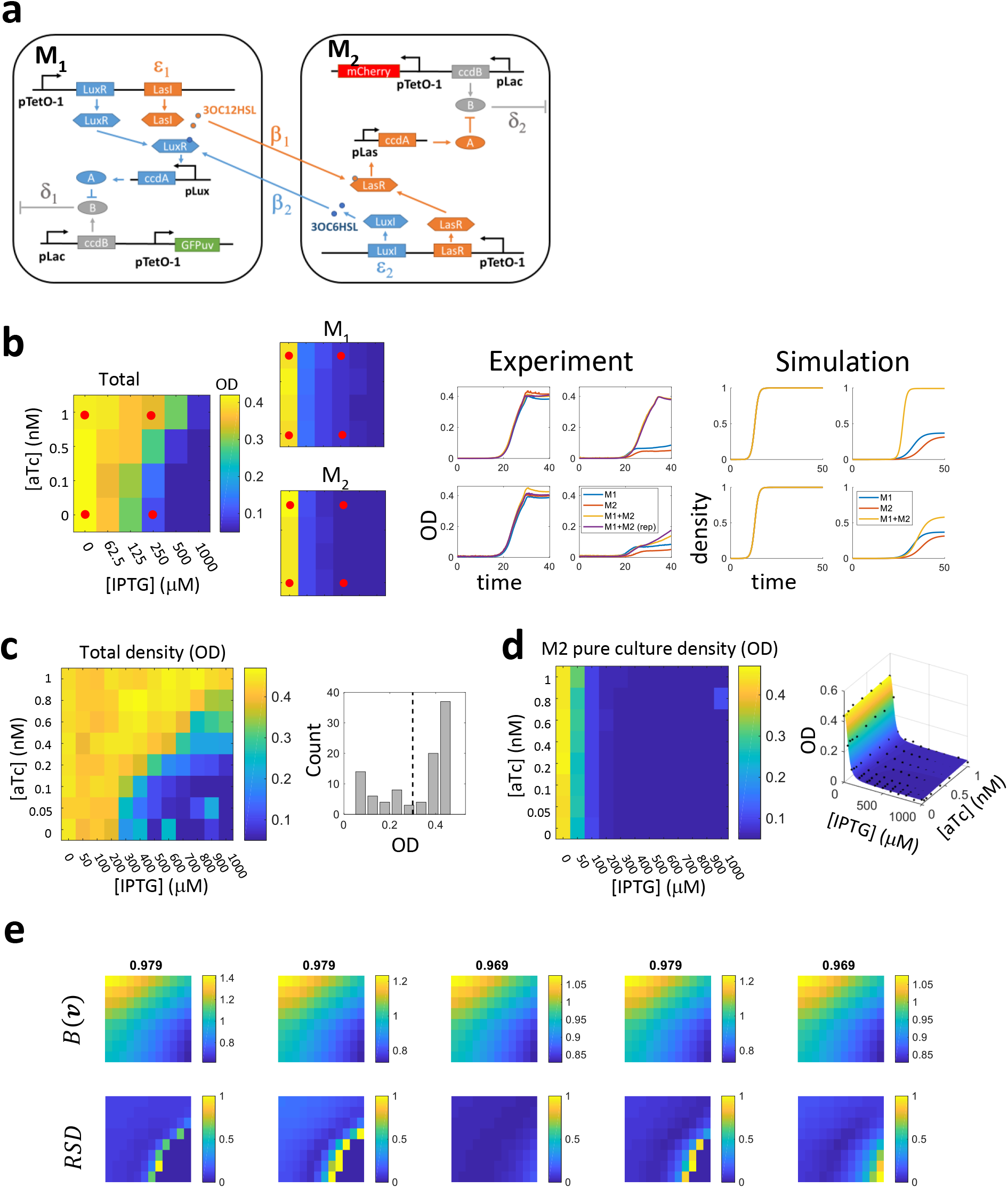
The QS-based mutualism system. **a**. The schematic of the synthetic gene circuit. *δ*_1_, *δ*_2_, *β*_1_, *β*_2_, *ε*_1_, and *ε*_2_ are indicated in the diagram to show the molecular mechanisms each term is primarily associated with. **b**. Verification of the basic system dynamics. The final OD values of coculture and monocultures are recorded. Monocultures of M_1_ and M_2_ are significantly suppressed by IPTG, while the cocultures exhibit synergistic growth. The total density of the cocultures increases with increasing [aTc]. The comparison between experimental and simulated temporal dynamics indicate that our model captures the basic mutualistic dynamics observed empirically. The four red dots in the heatmaps indicate the four experimental conditions of the simulated and experimental time courses. We used equations III.22 and 23 in the Supplementary Text for the simulation. The parameter values are: *δ*_1_([IPTG] = 0) = *δ*_2_([*IPTG*] = 0) = 0; *δ*_1_([IPTG] = 1000μM) = *0.63*; *δ*_2_([IPTG] = 1000μM) = 0.68; *β*_1_([aTc] = 0μM) = *β*_2_([aTc] = 0μM) = 2; *β*_1_([aTc] = 1nM) = *β*_2_([aTc] = 1nM) = 200; *ε*_1_ = *ε*_2_ = 1; *ρ* = 1 **c**. Using higher resolution of [aTc] and [IPTG] gradients, we obtained the total density of cocultures using OD measurements. At 32 hours, a bimodal distribution of final densities emerges with a trough at OD=0.3. **d**. Quantification of *δ* based on OD of M_2_ monoculture at 32 hours. The final OD is fitted using OD([*aTc*], [*IPTG*]) = 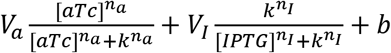 to reduce noise of the resulting *δ* measurements. *δ* is then calculated by *δ* = 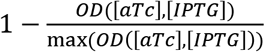. **e**. The top 5 *B*(***v***) and their corresponding relative standard deviation (*RSD*) (the axes are the same as heatmaps in panel c and d).

**Figure S7:**
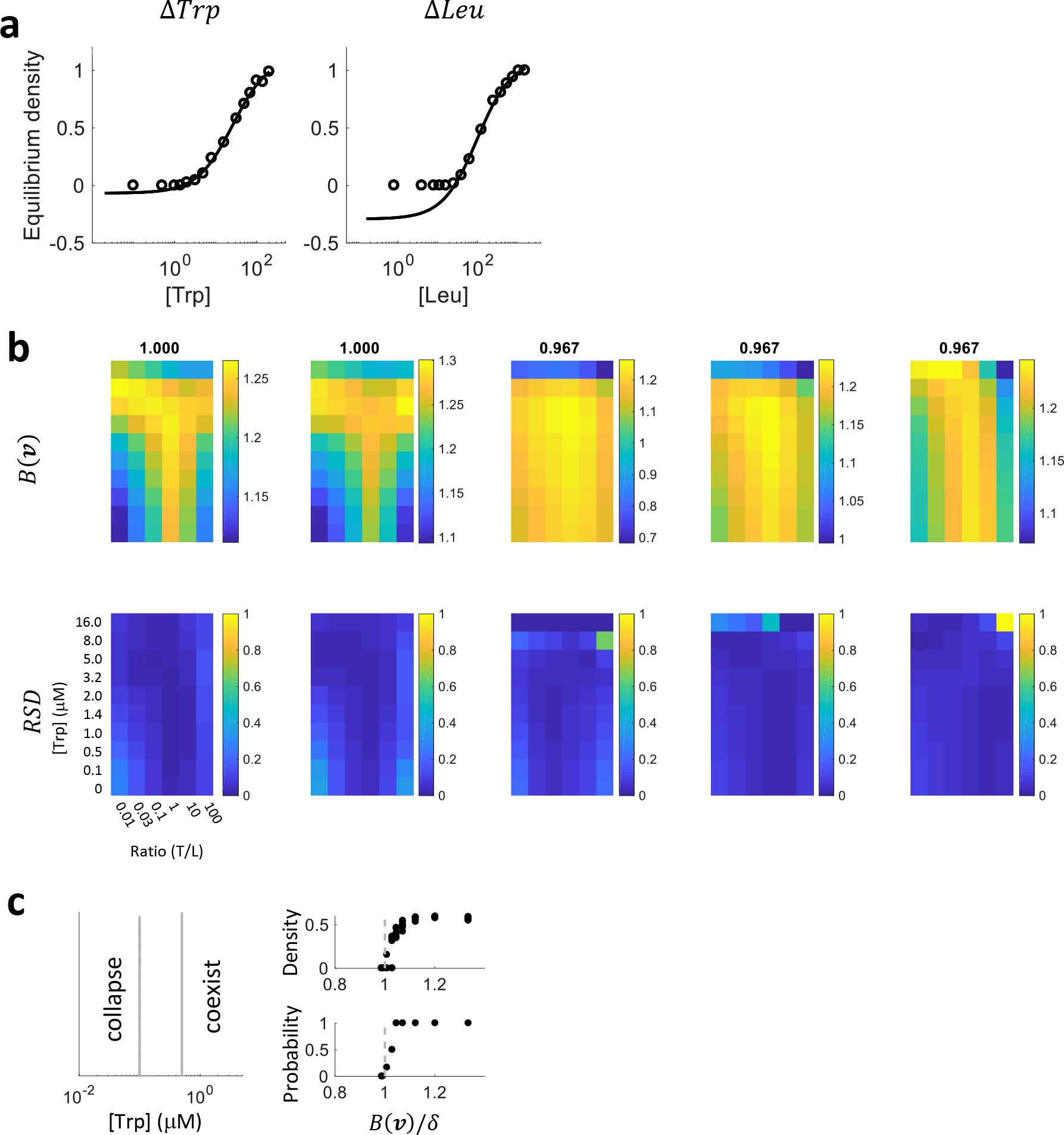
The yeast auxotroph system. **a**. The normalized equilibrium density of Δ*Trp* and Δ*Leu* were used to quantify *δ* for each strain. We only fitted density that is above the detection limit (10,000 cells/well). Since Δ*Leu* reaches below detection limit when [Leu] is relatively low, extrapolation is done to estimate the death rate at lower [Leu] (see Fig. S3a for the reasoning). As a result, *δ* of Δ*Leu* has a higher dynamic range than that of Δ*Trp*. **b**. 5 top *B*(***v***) show both consistency and discrepancy. All *B*(***v***) indicate an optimal initial density at an intermediate level. However, when [Trp] increase to 16 μM, *B*(***v***) can either increase or decrease. **c**. Our approach can also predict probability of coexistence. We excluded the ratio of initial density, leaving [Trp] to construct a one dimensional ***v***. The boundary between coexistence and collapse is between 0.1 μM tryptophan, 0.8 μM leucine and 0.5μM tryptophan, 4.0 μM leucine. To calibrate a *B*(***v***) for this system, we assume supplementing the two amino acids does not change the cooperation capability. This calibration demonstrates that our procedure can also predict probability of coexistence for experimental systems.

**Figure S8:**
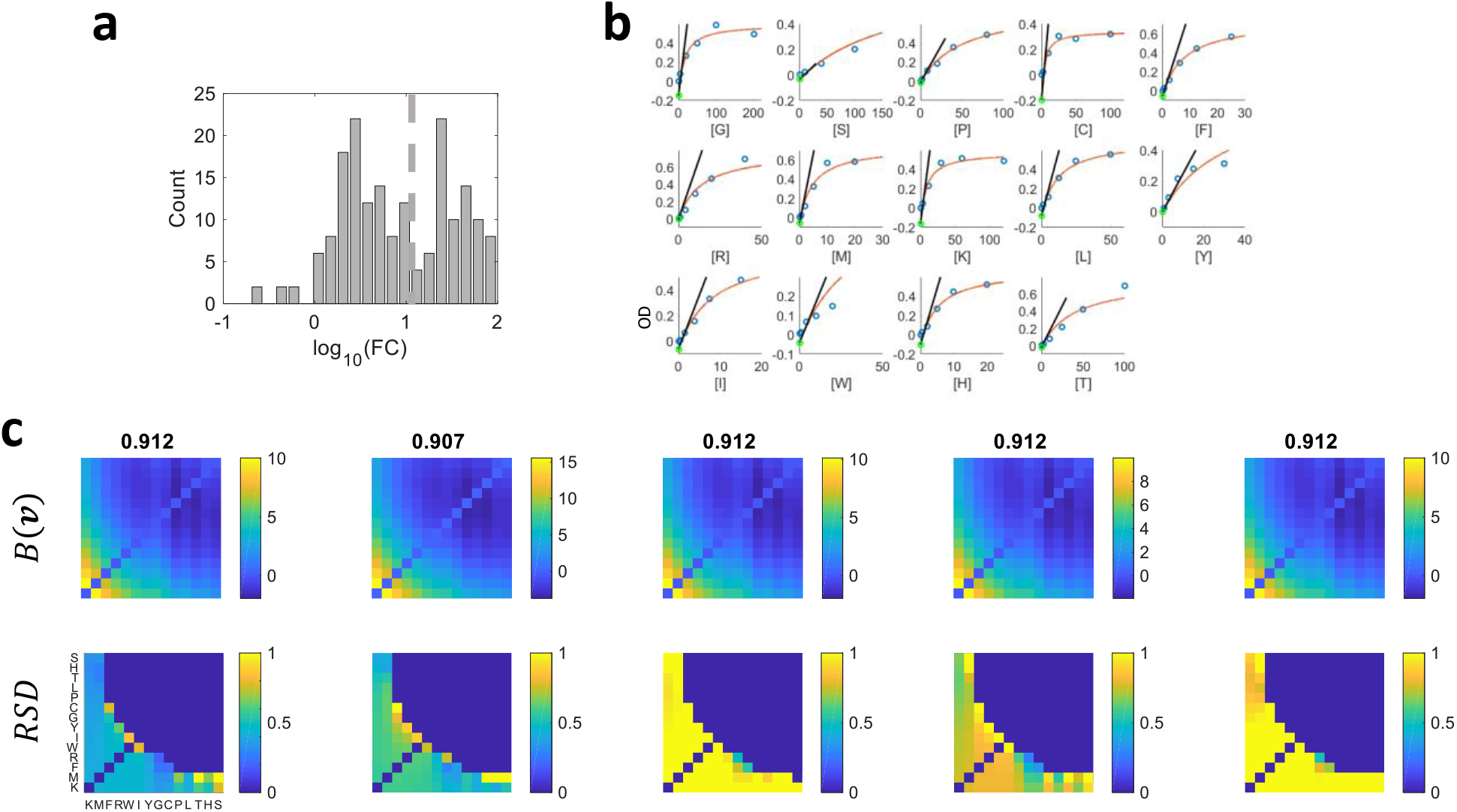
The 92 pairwise *e. Coli* mutualistic systems constructed using 14 auxotrophic strains. **a**. The logarithm of fold change (FC) of total final density for the 92 systems exhibit a bimodal distribution. The trough of the bimodal distribution corresponds to FC=10 (dashed grey line), which is used to classify coexistence and collapse. **b**. Quantification of *δ* for all 14 auxotrophs by fitting yield of the strain at different concentrations of their corresponding amino acid. Since none of the auxotrophs can grow without supplementary amino acid, *δ* ≥ 1 is expect for all auxotrophs at [AA]=0 (AA denotes the corresponding amino acid). Therefore, according to the reasoning demonstrated in Fig. S3, extrapolation of the data is required. We only fitted the data points that have OD>0.1, since OD<0.1 can be below the linear detection range of microplate readers. The data are fitted and extrapolated with Hill equations (red curves) or using a linear extrapolation of the fitted curves (the black trace). Green circles represent the values of OD of the fitted Hill equations at [AA]=0. *δ* is then calculated by 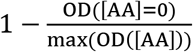 for all 14 strains using *OD*(*[AA]* = 0) obtained from either Hill equation or linear function. The two methods yield comparable results. We chose the *δ* calculated using the Hill equation for the calibration process. **c**. The top calibrated *B*(***v***) and their variability are indicated by relative standard deviation (*RSD*).

**Figure S9:**
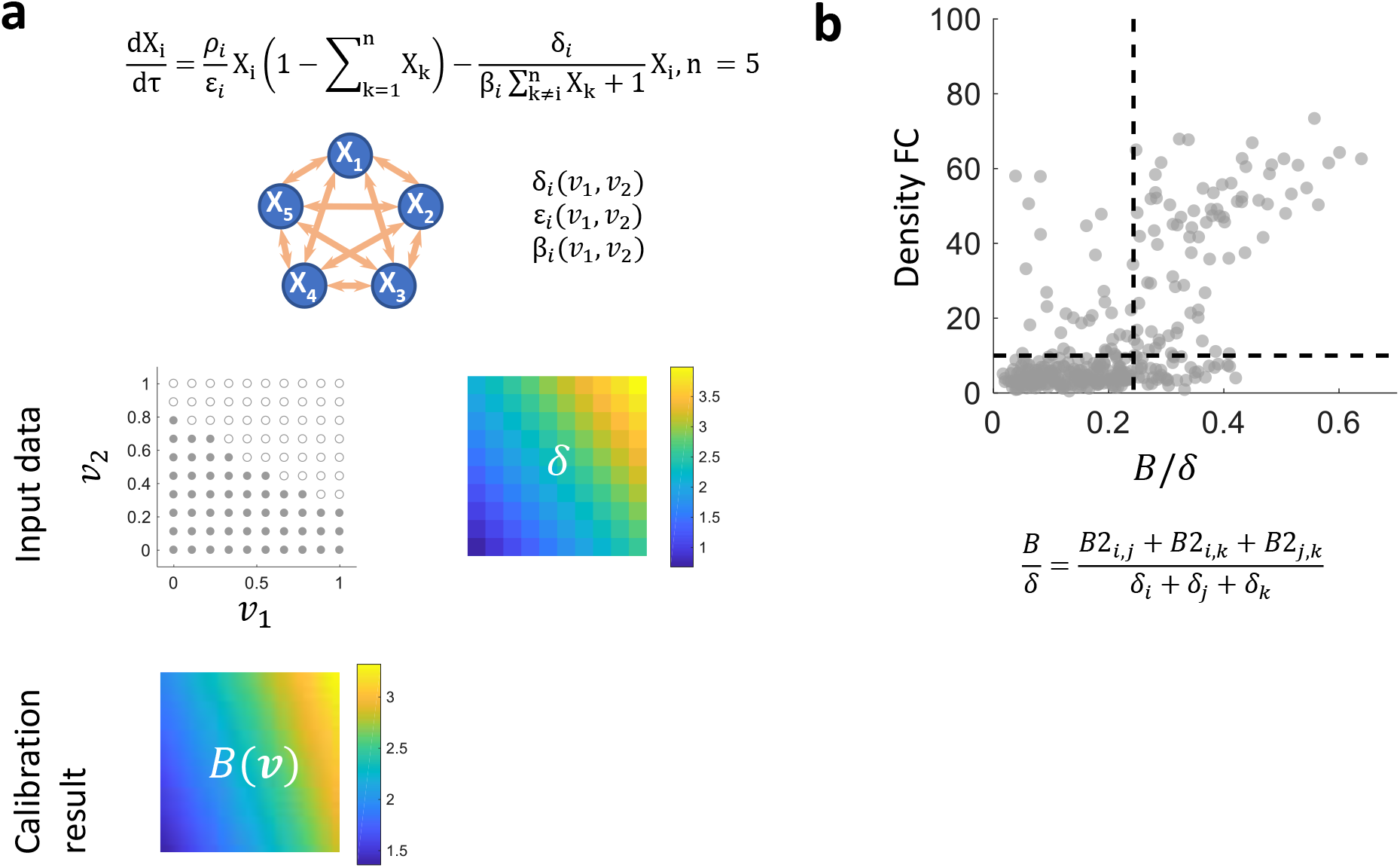
Mutualistic systems comprised of more than two partners. **a**. A mutualistic model with 5 partners. All the parameters in the model are linear functions of ***v***_1_ and ***v***_2_ and the parameters of the linear function are randomly generated. **b**. Constructing a predictive metric for 3-member systems from calibrated results of 2-member systems. We first normalized the range of calibrated *B* of 2-member systems (*B2*) to [0, 1]. The effective benefit of 3-member system is simply the average of *B2* for all underlying 2-member systems and stress is the average of *δ* for all 3 members. The subscripts of *B2* and *δ* represent the indices of the 3 members. We swept a threshold for *B/δ* between 0 and 1 to classify coexistence (FC≥10) versus collapse (FC< 10). The prediction accuracy reaches the maximum of *80.8%* at a threshold equals to 0.24.

**Figure S10:**
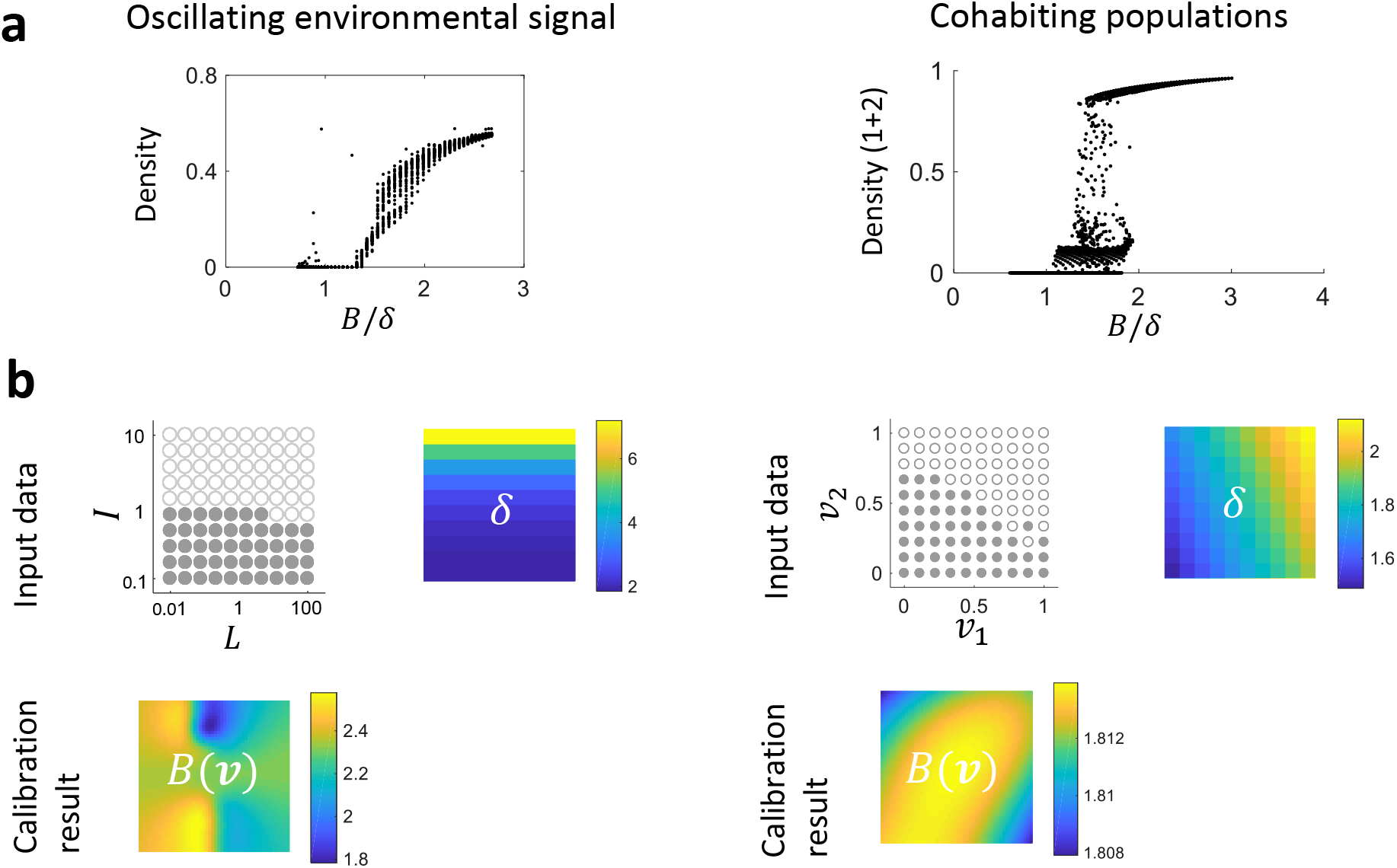
Mutualistic systems in dynamic environments. **a**. Two systems were analyzed: one system is a pairwise mutualism system that is modulated by oscillatory signals (the left panel) and the other system contains a pair of mutualists and 5 bystander populations that interact with the mutualists (the right panel). We verified that theoretically calculated *B/δ* roughly holds as a predictor for total density. The transition between coexistence and collapse, however, do not occur strictly at 1. Each dot represents a simulation result from one set of model parameters. **b**. The data used for the calibrations and the calibrated *B*(***v***). The two *B*(***v***)’s are used for the prediction plots shown in Fig. 4c, d.

**Movie S1**. This video demonstrates the relative position of the boundary surface *F(δ, **v***) = 0 and 5 observations in the schematic shown in Fig. 2b. Input data in Fig. 2b can be represented in a 3D space. The color of the dots and the z axis positions both indicate *δ* values. Coexistence is indicated by closed circles and collapse is indicated by open circles. The gray boundary surface separates coexistence and collapse. The surface is above observations that represent coexistence (*B* > *δ*) and below observations that represent collapse (*B* < *δ*). This surface can be directly interpreted as *B*(***v***) since *F(δ, **v***) = 0 ⟹ *F(*B*, **v***) = 0 ⟹ *B*(***v***)).

**Movie S2 – S4**. These 3 videos show the calibrated surface of *B*(***v***) relative to input data in 3D space. The blue surfaces are the boundary *F(δ, **v***) = 0 that separate coexistence and collapse. The surface is also equivalent to *B*(***v***). The indices on x and y axes in video 4 correspond to the same strain orders in Fig. 3j.

MATLAB code and movie S1 – S4 can be found using the following Dropbox link: https://www.dropbox.com/sh/c65vqo0vw4sxb6k/AAC4QpRXs9KyiCZV1RjMaPfqa?dl=0

